# Investigation of the molecular-genetic basis for courtship differences between *Drosophila melanogaster*, *Drosophila simulans* and their hybrids

**DOI:** 10.1101/2025.08.05.668758

**Authors:** Colleen M. Palmateer, Kelsey Claire Morris, Scott Nolting, Jackson Mast, Michelle N. Arbeitman

## Abstract

Understanding the mechanisms driving behavioral evolution enhances our knowledge of speciation and how behavioral potentials are genetically encoded. Male courtship behaviors, which evolve rapidly, are critical for pre-mating isolation. To investigate the genetic basis of species-specific courtship, we generated hybrid males by crossing two *Drosophila melanogaster* strains with *D. simulans Lhr* males. This design allowed us to assess how genetic background and female species identity influence male courtship behavior. In both single-pair and mate-choice assays, hybrid males displayed behavioral plasticity, adjusting their courtship strategies based on the female species. The maternal *D. melanogaster* strain significantly shaped hybrid behavioral repertoires. To identify the neural correlates, we examined *fruitless* (*fru*)-expressing neurons in hybrids. Their projection patterns resembled those in *D. melanogaster*, indicating conserved circuit architecture. Using CUT&Tag, we identified Fru^M^ target genes in both species, revealing conserved core and species-specific Fru^M^ targets. Focusing on chemosensory receptors with *D. melanogaster*-specific Fru^M^ binding, we conducted a genetic screen in *D. melanogaster*, silencing neurons co-expressing *fru P1* and specific receptor genes. Courtship preference assays identified three additional olfactory receptor neuron subtypes that modulate species-specific behavior. Using *trans-Tango*, we mapped second-order projection neurons of these subtypes, revealing their targets in higher-order brain centers. This study uncovers new molecular and neural mechanisms underlying the specification and evolution of courtship behavior, highlighting how genetic and sensory inputs shape species-specific behavioral outcomes.

**Article Summary:** We examined how genes and circuits shape courtship behaviors in *Drosophila melanogaster* and *Drosophila simulans*. Hybrid males adjust their behavior based on the species of female, suggesting flexible courtship. We focused on the *fruitless* gene, which controls male courtship. *fru* neurons in hybrids closely resembled those in *Drosophila melanogaster*. We identified both shared and species-specific Fru^M^ target genes. A genetic screen of odorant receptor neurons revealed populations that help males distinguish females. Neuroanatomical mapping shows the odor information is relayed to different higher order brain regions. The results uncover molecular and neural circuit mechanisms underlying species differences in behaviors.

## Introduction

Elucidating the mechanisms that drive behavioral evolution is important to both further our understanding of speciation and to provide insights into how the potential for behaviors are specified. Male courtship behaviors are rapidly evolving traits that play a significant role in pre-mating isolation across species (reviewed in ANHOLT *et al*. 2020). The model organism *Drosophila melanogaster* is a powerful system for understanding the molecular-genetic basis of behavioral evolution, and has provided insights into how molecular changes direct differences in neuronal connectivity, gene expression, and physiology to produce species-specific reproductive behaviors (reviewed in ANHOLT *et al*. 2020; SATO *et al*. 2020). Core elements of courtship activity, such as wing vibration, licking, and circling are common to all species within the *Drosophila* genus (SPIETH 1952; SPIETH 1974), with quantitative differences across behavioral elements distinguishing closely related species, while more distantly related species have larger qualitative behavioral differences. Courtship behaviors across the *melanogaster* species subgroup, which consists of nine species (for history of discovery see: DAVID *et al*. 2007), mostly have quantitative differences in behavioral elements, with some changes in the courtship elements (COBB *et al*. 1985; COBB *et al*. 1986). This study focuses on species differences for courtship behavior between *D. melanogaster* and *D. simulans* (**Figure 1**), two closely related species, which exhibit distinct courtship behavior and mate preferences.

**Figure 1.**
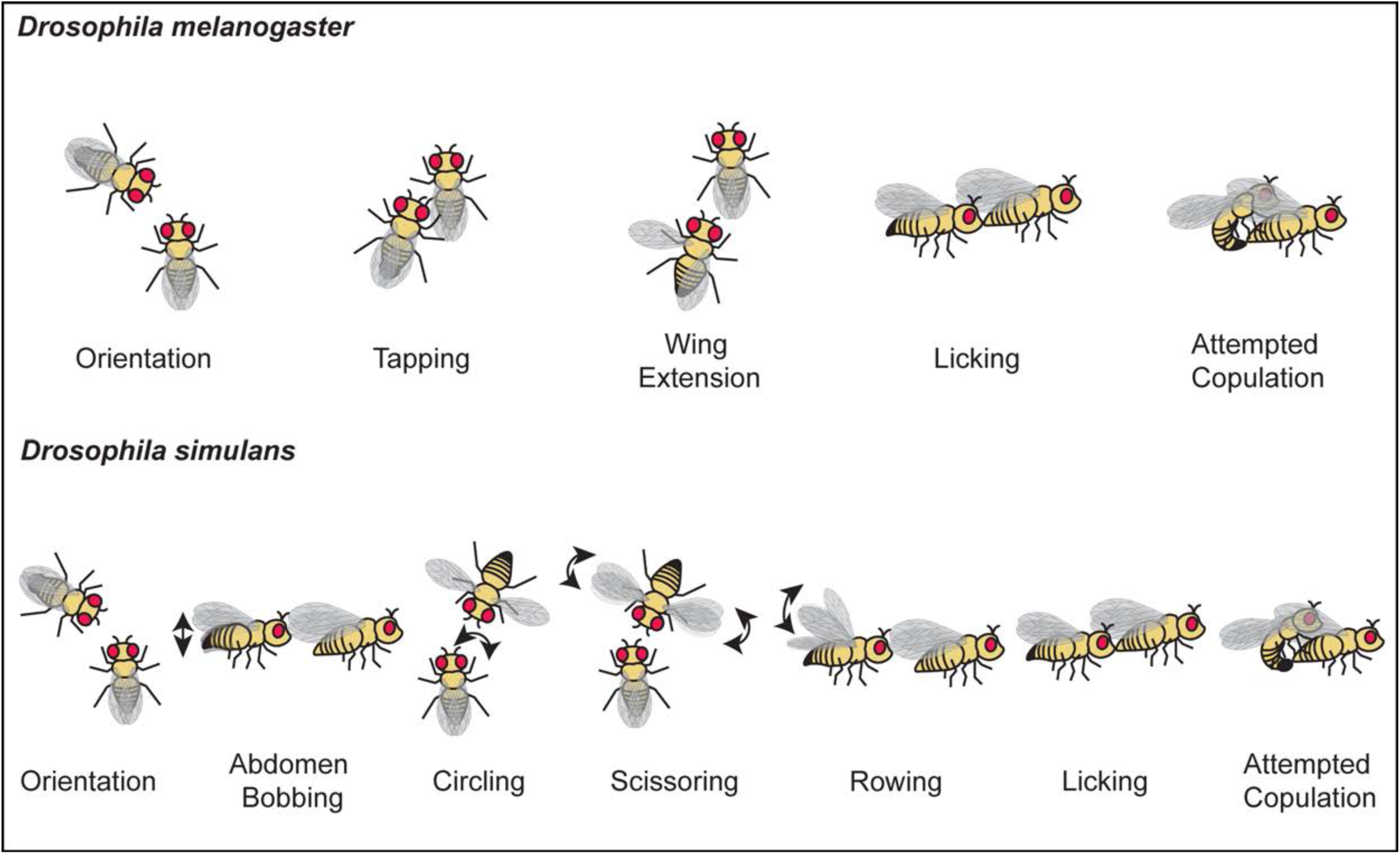
Courtship behaviors of *D. melanogaster* and *D. simulans*. The *D. melanogaster* male initiates courtship by orienting towards the female, he then taps her abdomen with his foreleg, extends one wing to sing a species-specific courtship song, extends his proboscis to lick her genitalia, and attempts copulation. In contrast, the *D. simulans* males display additional behaviors not commonly observed in *D. melanogaster* males. These include abdomen bobbing, circling the female while flaring out his wings, wing scissoring (flaring out wings and vibrating them) and wing rowing (extending one or two wings, raising them up and then rowing them back to their body without vibration).

*D. melanogaster* and *D. simulans* diverged approximately 3.5-5 million years ago (TAMURA *et al*. 2004), and are sympatric (reviewed in JEZOVIT *et al*. 2017). In both species, male courtship behavior involves a series of stereotyped repeated elements that relies on several sensory modalities, including vision, taste, olfaction, sound, and touch (reviewed in GREENSPAN AND FERVEUR 2000; ANHOLT *et al*. 2020). Courtship behavior begins with the male orienting toward the female and tapping her body with a foreleg. If the female is walking, the male will pursue her by following while performing wing displays, singing a species-specific courtship song, and circling her, often combining these behaviors simultaneously (e.g. following and wing extensions or song and circling).

In *D. melanogaster*, males will first orient themselves toward any portion of the female target, tap her abdomen with one foreleg, and pursue her if she is mobile (**Figure 1**) (reviewed in GREENSPAN AND FERVEUR 2000). The male will unilaterally extend and vibrate one wing, at 50° or 90° angle, to produce a species-specific courtship song, that can be interrupted by bouts of genital licking, and copulation attempts (**Figure 1**). If the female is receptive, this sequence will ultimately lead to copulation. There have been reports of wing scissoring and abdominal tremulations as a part of the *D. melanogaster* courtship sequence (COBB *et al*. 1985; COBB *et al*. 1986; FABRE *et al*. 2012).

*D. simulans* males exhibit a similar courtship sequence, initially orienting toward the female’s head or genitalia at close range and following her, as in *D. melanogaster* courtship (**Figure 1**) (STURTEVANT 1915). They also perform a distinctive behavior, moving in an arc in front of the female in a circling motion, often repeating this several times in succession, typically accompanied by wing scissoring (**Figure 1**). Both wing vibrations and wing scissoring are characterized by the extension of one wing, or both in the case of scissoring, at approximately 45° degrees, with a slight elevation (**Figure 1**). The wing scissoring motion has been noted to surge in wing separation and speed of wing closure. Wing rowing will also be performed by one or both wings by extension of 45-50° and a raise to 30°, without visible vibration (COBB *et al*. 1985). These behaviors are often accompanied by abdomen bobbing, which can occur during orientation or while the male performs wing vibration and rowing behaviors. As in *D. melanogaster,* genital licking, attempted copulation, and copulation are part of courtship (**Figure 1**) (STURTEVANT 1915).

Despite some behavioral similarities, these two *Drosophila* species differ in their mate preferences. *D. melanogaster* males will readily court *D. simulans* females, whereas *D. simulans* males show a strong preference for conspecific females. These preferences are largely driven by chemosensory perception of species-specific cuticular hydrocarbons (CHCs) (reviewed in JALLON 1984; FERVEUR 1997; BONTONOU AND WICKER-THOMAS 2014) (AVERHOFF AND RICHARDSON 1974; JALLON AND DAVID 1987; BILLETER *et al*. 2009; COMBS *et al*. 2018; SHAHANDEH *et al*. 2018). The CHCs on female *Drosophila* act as pheromones, detected by the gustatory and olfactory system, facilitating species recognition and mate selection (reviewed in SATO AND YAMAMOTO 2020) (FERVEUR AND SUREAU 1996; BILLETER *et al*. 2009; PARDY *et al*. 2019). *D. melanogaster* females primarily produce 7,11 heptacosadiene (7,11-HD) and *D. simulans* females primarily produce 7-tricosene (7T) (reviewed in SATO *et al*. 2020) (JALLON 1984; BILLETER *et al*. 2009). Notably, 7,11-HD is attractive to *D. melanogaster* males but acts as a repulsive cue for *D. simulans* males, contributing to reproductive isolation between the species.

Over the last decade, there has been significant progress in understanding the neural circuit basis for courtship and conspecific partner preference in *Drosophila* (reviewed in ANHOLT *et al*. 2020; SATO *et al*. 2020). Much of this work has focused on a class of neurons that express male-specific Fru (Fru^M^) transcription factors, encoded by *fruitless* (*fru*; *fru P1* transcript class encodes Fru^M^) (for example see TOOTOONIAN *et al*. 2012; FAN *et al*. 2013; TANAKA *et al*. 2017; SEEHOLZER *et al*. 2018; BAKER *et al*. 2024; COLEMAN *et al*. 2024) (reviewed in CLYNEN *et al*. 2011; PENG *et al*. 2021; SATO AND YAMAMOTO 2023). In *D. melanogaster*, *fru P1*-expressing neurons have been shown to underlie nearly all aspects of male courtship behaviors. *fru P1* neurons are estimated to be 2-5% of all neurons, and reside in both peripheral sensory neurons, and throughout the central nervous system (reviewed in AUER AND BENTON 2016; ANHOLT *et al*. 2020; PENG *et al*. 2021) (MANOLI *et al*. 2005; STOCKINGER *et al*. 2005).

In *D. melanogaster* males, 7,11-HD is perceived by gustatory receptor neurons in the foreleg tarsals that express ion channel genes *pickpocket 23* (*ppk23*) and *pickpocket 25* (*ppk25*) (LU *et al*. 2012; THISTLE *et al*. 2012a; TODA *et al*. 2012; LU *et al*. 2014). These neurons relay signals to *fru P1*-expressing vAB3 and non-*fru P1* PPN1 ascending interneurons (CLOWNEY *et al*. 2015; KALLMAN *et al*. 2015), which then signal to male-specific *fru P1*-expressing neurons called P1, located in the protocerebrum (KIMURA *et al*. 2005; CACHERO *et al*. 2010; YU *et al*. 2010), whose activation has been shown to promote male courtship behaviors (KOHATSU *et al*. 2011; VON PHILIPSBORN *et al*. 2011; PAN *et al*. 2012; INAGAKI *et al*. 2014; ISHII *et al*. 2020). The vAB3 and PPN1 ascending interneurons also send information to mAL neurons (KIMURA *et al*. 2005; CACHERO *et al*. 2010; YU *et al*. 2010; ITO *et al*. 2012), which are sexually dimorphic *fru P1*-expressing GABAergic interneurons (KOGANEZAWA *et al*. 2010; KALLMAN *et al*. 2015; reviewed in AUER AND BENTON 2016). When activated, mAL neurons inhibit P1 neurons to inhibit courtship pathways (CLOWNEY *et al*. 2015; KALLMAN *et al*. 2015). Thus, 7,11-HD perception drives courtship through a balance of excitatory (via P1) and inhibitory (via mAL) pathways (CLOWNEY *et al*. 2015; KALLMAN *et al*. 2015). Additional pathways have been identified for other pheromones that mediate courtship behaviors through gustatory and olfactory pathways (for examples see THISTLE *et al*. 2012b; LEBRETON *et al*. 2017; VERNIER *et al*. 2023; LUO *et al*. 2024).

Recent research examined the neural basis of courtship inhibition by 7,11-HD in *D. simulans* males (SEEHOLZER *et al*. 2018). They evaluated *fru P1*-expressing neurons in *D. simulans* by generating transgenic strains that label these neurons. They found the morphology of the *fru P1*-expressing neurons was largely homologous to *D. melanogaster* and that these neurons also control male courtship. This suggests that species-specific courtship behaviors arise from subtle changes in neuronal architecture, or physiology. They found that 7,11-HD inhibits courtship in *D. simulans* males via enhanced mAL-mediated inhibition of the P1 neurons through the vAB3 circuit (SEEHOLZER *et al*. 2018).

Speciation often involves pre- and post-mating isolation barriers that reduce gene flow (COYNE AND ORR 2004), including behavioral isolation, hybrid inviability and sterility. Many *melanogaster* subgroup species can hybridize (LEE AND WATANABE 1987; CANDE *et al*. 2012), though F_1_ progeny are often sterile, or produce viable progeny of only one sex (STURTEVANT 1921) (reviewed in BARBASH 2010). These hybrid incompatibilities typically result from divergent gene interactions (for example see: CATTANI AND PRESGRAVES 2009; WEI *et al*. 2014), with many mapped to specific chromosomes, and some causative genes identified (HOLLOCHER and Wu 1996; True *et al*. 1996; Sawamura and Yamamoto 1997; Presgraves *et al*. 2003; Tao *et al*. 2003; Presgraves and Masly 2007; Cuykendall *et al*. 2014; Cooper *et al*. 2019).

Many of these incompatibilities are linked to protein-coding genes (TING *et al*. 1998; BARBASH *et al*. 2003; PRESGRAVES *et al*. 2003; BRIDEAU *et al*. 2006; PHADNIS *et al*. 2015; TANG AND PRESGRAVES 2015), though, repeat regions (SAWAMURA AND YAMAMOTO 1997; CATTANI AND PRESGRAVES 2012), and translocations (MASLY *et al*. 2006) have also been identified as causative.

Crosses between *D. melanogaster* females to *D. simulans* males produce no F_1_ sons, and the reverse cross yields no daughters (STURTEVANT 1920). A natural *D. simulans* strain, *Lhr^1^*, was found to rescue this hybrid lethality (WATANABE 1979). Lhr interacts with HP1, a heterochromatin-associated protein, in *D. melanogaster* and it has been hypothesized that this is also the case in *D. simulans* (BRIDEAU *et al*. 2006). The *Lhr*^1^ *D. simulans* mutant strain contains a ∼4kb insertion that generates a loss-of-function allele that restores dosage compensation on the X chromosome (BRIDEAU *et al*. 2006; CHATTERJEE *et al*. 2007). This enables the production of viable hybrid progeny of both sexes. These hybrids, containing genetic material from both species, offer a unique system to study the genetic basis of behavior. Their courtship patterns, songs and mate preferences have been extensively analyzed (VON SCHILCHER AND MANNING 1975; WOOD AND RINGO 1980; WOOD *et al*. 1980; SHAHANDEH *et al*. 2020).

To examine the genetic basis underlying species differences in courtship behavior, we generated hybrid males by crossing two strains of *D. melanogaster* females with *D. simulans Lhr* males. This design enabled us to assess how genetic background influences the expression of species-specific courtship behaviors and whether these behaviors are specified in a quantitative manner. Additionally, this approach allowed us to evaluate the role of the female courtship target in shaping male behavioral responses. In both single-pair and mate-choice preference assays, hybrid males exhibited behavioral plasticity, modulating their courtship strategies depending on the species of the female. Notably, the *D. melanogaster* maternal strain significantly influenced the behavioral repertoire of the hybrids. To investigate potential neural correlates of these behavioral differences, we examined *fru*-expressing neuronal subsets in hybrid males. The projection patterns observed were highly similar to those in *D. melanogaster*, suggesting conservation of circuit architecture despite the hybrid genetic background. To gain insight into the molecular mechanisms driving species-specific behaviors, we identified male-specific Fruitless (Fru^M^) targets in *D. melanogaster* and *D. simulans*, using the CUT&Tag approach (KAYA-OKUR *et al*. 2019). This analysis revealed a core set of conserved Fru^M^ target genes, as well as species-specific targets. Given the role of pheromone signaling in pre-mating isolation, we focused on chemosensory receptors exhibiting *D. melanogaster*-specific Fru^M^ binding.

Several such receptors were identified. To functionally assess the role of these receptors, we performed a genetic screen in *D. melanogaster,* using a genetic intersectional approach to silence neurons co-expressing *fru P1* and specific chemosensory receptor genes. In courtship preference assays, with both *D. melanogaster* and *D. simulans* females, we identified three additional olfactory receptor neuron subtypes that co-express *fru P1* and modulate species-specific courtship behavior (*Or42b*-, *Or43a*-, and *Or43b*-expressing neurons). We used the *trans*-Tango approach to evaluate their second-order olfactory projection neurons, revealing differences in their targets in higher-order processing centers and suboesophageal ganglion (TALAY *et al*. 2017). This study identifies new molecular mechanisms underlying how the potential for behavior is established and provides insights into the substrates for evolution of these behaviors.

## Materials and methods

### Drosophila husbandry, hybrid generation, and strains used in this study

The model systems used in this study are *Drosophila melanogaster* (*D. melanogaster*) and *Drosophila simulans* (*D. simulans*). All strains, unless otherwise indicated, are grown in humidified incubators at 25°C on a 12-hour light and 12-hour dark cycle. The laboratory food media composition is: 33 L H2O, 237 g agar, 825 g dried deactivated yeast, 1560 g cornmeal, 3300 g dextrose, 52.5 g Tegosept in 270 ml 95% ethanol and 60 ml propionic acid.

A full list of stocks used in this study is provided (**Supplementary Table 1**). Two different crosses were performed to generate *D.melanogaster* and *D. simulans* hybrid males to examine behavior. (**1**) One cross was between Berlin virgin females and *D. simulans Lhr^1^* males. The F_1_ males from this cross are called Hybrid_1. (**2**) The other cross was between virgins of a *D. melanogaster w^1118^* strain (from Well Genetics in Taiwan) that had the *white* gene introgressed back into the background, by five generations of backcrossing (hereafter called *w+*WG). This *D. melanogaster* strain was chosen as virgins would readily mate with *D. simulans Lhr^1^* males. The F_1_ males from this cross are called Hybrid_2. Additional hybrids were generated to examine neuronal anatomy by confocal imaging. To generate F_1_ adult hybrid progeny of both sexes for all genotypes used in this study, we crossed *D. melanogaster* females to *Lhr^1^ D. simulans* males. To set up hybrid crosses, we collected young 0-6 hours post-eclosion *Lhr^1^D.simulans* males and aged them at least 4-7 days. 10-15 of these aged *Lhr^1^ D. simulans* males were combined with at least 10 young *D. melanogaster* females (0-6 hours post-eclosion). We added a piece of rolled and H_2_O moistened Kimwipe into the food, in a bridge-like configuration. For some crosses, the cotton plug was pressed into the vial to just above this Kimwipe bridge, resulting in about 2cm of space.

### Male courtship assays with hybrid males

For male courtship assays, all male flies were collected within 0-6 hours post-eclosion and aged individually for 4-10 days. *D. melanogaster* and *D. simulans* virgin female flies were collected and housed in groups of ∼10 and aged for 4-10 days. For single-pair courtship assays, a single male is paired with either a single virgin *D. melanogaster* or *D. simulans* female. For the preference courtship assays a single male is paired with a virgin *D. melanogaster* and a virgin *D. simulans* female. The assays were performed using a 10mm circular chamber set on a 25°C temperature block, 5-9 hours after lights-on. Recordings were analyzed using Observer® XT software (Noldus). The n and statistical tests are provided in Supplementary Table 2.

For the courtship assays, we calculated indices by dividing the time the experimental male fly spent performing each behavior toward the target female by the total observation time, or until successful mating. The Courtship Index (CI) is the amount of time the male was following the female. We additionally quantified specific courtship behavior elements including Wing Extension Index (WEI), Wing Scissoring Index (WSI), Abdomen Bobbing Index (ABI), and Circling Index. Counts of attempted copulations and percentage of copulation successes were also analyzed.

The preference indices were calculated as the amount of time the male spent performing a behavior toward the *D. melanogaster* female subtracted from the amount of time spent courting the *D. simulans* female divided by the total behavior performance time. For preference calculations, males who did not perform the respective behavior (following, wing extension, or wing scissoring) were removed from the analysis, as they would have no preference. GraphPad Prism 9.3.0 was used to conduct statistical analyses. The statistical tests and n are indicated in the figure legends and summary statistics table (**Supplementary Table 2**)

### Immunohistochemistry

Brains and ventral nerve cords were dissected from either 0-24 hour or 5–10-day old adults, as indicated. Tissues were dissected in 1x PBS (PBS; 140 mM NaCl, 10 mM phosphate buffer, and 3 mM KCl, pH 7.4) and fixed for 20 +/- 2 minutes in 4% PFA (Electron Microscopy Sciences, product 15713), made in 1x PBS at room temperature. The tissue next underwent 4x one-minute 1x PBS washes and were then washed once with TNT (0.1M Tris-HCL, 0.3M NaCl, 0.5% Triton X-100) for 15 min and then 2x more times for 5 minutes per wash. The tissue was blocked in Image-iT FX Signal Enhancer (Thermo Fisher) for 25 minutes then washed 2x with TNT for 5 minutes each. Tissue was incubated in primary antibody diluted in TNT overnight at 4°C. After removal of primary antibody, the tissue was washed 6x with TNT for 5 minutes each wash. The tissue was then incubated in TNT with secondary antibody for two hours at room temperature or overnight at 4°C. Finally, secondary antibody was removed, and the tissue was washed 6x for 5 minutes in TNT, before being mounted on glass slides with Secureseal Image Spacers (Electron Microscopy Services) in Vectashield mounting medium (Vector Laboratories, H-1000) and covered with a no. 1.5 glass coverslip. Confocal microscopy was performed on a Zeiss LSM 700 or 900 system (Odorant and Gustatory Receptor studies), with a Zeiss Plan-Apochromat 20x/0.8 M27 objective.

For the *trans*-Tango experiment, crosses were set up at 25°C. The F_1_ progeny were collected, transferred to 18°C and aged for 14-19 days. Dissections were performed as above, but primary antibody incubations were longer, and ranged from 2-4 full days at 4°C.

The following primary antibodies were used for immunohistochemistry: rat anti-Fru^M^ (1:200; (SANDERS AND ARBEITMAN 2008)), mouse anti-nc82 (1∶20; Developmental Studies Hybridoma Bank), mouse anti-FLAG (1:500; Sigma, F1804; for smGDP), chicken anti-myc (1:1000; Invitrogen, A21281; for smGDP), rabbit anti-HA-tag (1:300; Cell Signaling, C29F4; for MCFO and trans-Tango), mouse anti-bruchpilot (1:20, DSHB, nc82), chicken anti-V5 tag 550 (1:500; Novus Biologicals, NB600-379R; for MCFO), mouse anti-GFP (1:100; DSHB GFD-4C9-b; for *trans*-Tango). The secondary antibodies used were Alexa Fluor goat anti-rat 488 (1:1000; for Fru^M^), goat anti-mouse 633 (1:500; for FLAG), goat anti-chicken 546 (1:1000; Invitrogen, A11040; for myc), rabbit α-GFP Alexa Fluor 488 (1:600; Invitrogen, A21311), goat anti-rabbit 488 (, 1:500; Invitrogen, 32731; for MCFO), goat anti-mouse 633 (1:500; Invitrogen, A21071, for MCFO), and goat anti-mouse 488 (1:500; Invitrogen, A110001; *trans*-Tango).

### Image registration

A *D. melanogaster*/hybrid male standard template brain and ventral nerve cord (interspecies) was generated with the Computational Morphometry Toolkit (CMTK, www.nitrc.org/projects/cmtk) (ROHLFING AND MAURER 2003; ROHLFING 2012), in FIJI using previously described parameters (CACHERO *et al*. 2010). The interspecies templates were generated by averaging images with the highest quality anti-nc82 staining. This resulted in 7 *D. melanogaster* brains and 10 hybrid brains averaged for the interspecies brain template. The interspecies male ventral nerve cord template was generated by averaging 28 *D. melanogaster* ventral nerve cords and 25 hybrid ventral nerve cords. Confocal stacks of the *fru P1* neuronal subsets from only the highest quality images obtained were registered, where anti-nc82 staining, a neuropil marker, was uniform and all brain and ventral nerve cord structures were present and appropriately positioned, resulting in a lower number of images going into this analysis than those obtained overall. *D. melanogaster* and hybrid males images were registered to the standard brain and ventral nerve cord templates, using linear registration and non-rigid warping based on the anti-nc82 staining channel (OSTROVSKY *et al*. 2013). Scripts to accomplish this registration were created using the GUI implementation of CMTK in FIJI.

### CUT&Tag to identify Fru^M^ target genes

Males used in CUT&Tag were 15-24 hours old. To collect these males, bottles with *D. melanogaster* and *D. simulans* were cleared at 10:00 AM, flies were collected at 5:00 PM, and aged overnight to allow for recovery from CO_2_ anesthesia. The male flies were stored individually in small vials with food, in a 25°C incubator. The next day, brain dissections started at 8:00 AM and ended at 10:00 AM. Ten brains were used for each CUT&Tag sequencing library. The CUT&Tag procedure and sequencing library preparation protocol was previously described, with some modifications (AHMAD 2020). After dissection, brains were fixed in 1% paraformaldehyde/1X PBS for 1 minute, then transferred to 1X PBS. Brains were washed two additional times for 5 minutes per wash. The rat polyclonal anti-Fru^M^ antibody was used at a 1:50 dilution (SANDERS AND ARBEITMAN 2008). The secondary rabbit anti-rat IgG (abcam ab6703) was used at 1:100 dilution. For control libraries, the brains were only incubated in secondary antibody. DNA recovery was performed as previously described (KAYA-OKUR *et al*. 2019). Samples underwent 21 cycles of PCR and were purified by agencourt bead purification before sequencing (AHMAD 2020). All libraries were sequenced on an Illumina NovaSeq 6000 in the Florida State University Translational Core Laboratory.

The CUT&Tag sequencing reads were pre-processed to remove Illumina adaptor sequences using BBduk (ktrim=r k=23 mink=11 hdist=1 tpe tbo (BUSHNELL *et al*. 2017) (https://sourceforge.net/projects/bbmap/). Reads were mapped and processed using the guidance from the CUT&Tag Data Processing and Analysis Tutorial (HENIKOFF *et al*. 2020; ZHENG Y 2020). A summary of these steps with the commands utilized are as follows. The adaptor trimmed reads were aligned to the *D*. *melanogaster* genome (BDGP6) genome assembly, using Bowtie2 (--local --very-sensitive --no-mixed --no-discordant --phred33 -I 10 -X 700 -p 8 -x /; v. 2.2.5) (LANGMEAD AND SALZBERG 2012). The mapped reads then underwent a series of file format conversion steps to prepare for SEACR peak calling. The mapped read files, in the .sam format, were converted to .bam files using Samtools view command (view -bS -F 0×04) (DANECEK *et al*. 2021). These .bam files were sorted using the Samtools sort command (sort -n - O bam -@ 12). The sorted .bam files were converted to .bed files using bedtools bamtobed command (bamtobed -bedpe -i). These files were filtered to keep read pairs on the same chromosome with a fragment length <1000bp (awk ‘$1==$4 && $6-$2 < 1000 {print $0}) and the fragment related columns were extracted using ‘cut -f 1,2,6’. These .bed files were converted to .bedgraph files for peak calling using bedtools (genomecov -bg -i). Peaks were called using SEACR (MEERS *et al*. 2019), using the ‘norm stringent’ parameters. The SEACR peaks for each replicate were relative to the species-specific IgG control data. The SEACR peaks were annotated with the default parameters in PAVIS, using the *D. melanogaster* FlyBase R6.05 genome release (HUANG *et al*. 2013; OZTURK-COLAK *et al*. 2024).

To identify Fru^M^ target genes for each species, we found genes with SEACR peaks in both data sets, resulting in 4,115 genes and 5,097 genes, identified in *D. melanogaster* and *D. simulans*, respectively (hereafter called *D. melanogaster* and *D. simulans* data sets; **Supplementary Table 3**). We found 2,874 genes with peaks in both the *D. melanogaster* and *D. simulans* datasets. We found 1,541 genes with peaks in only the *D. melanogaster* data set. We found 2,223 genes with peaks in only the *D. simulans* data set. We downloaded the list of genes that are annotated as “Chemoreceptor Genes” and “Ion Channel” from Flybase (OZTURK-COLAK *et al*. 2024), and identified the overlapping genes from the *D. melanogaster* and *D. simulans* data sets.

We performed enrichment analyses on the gene lists, examining overlaps with previous studies examining Fru^M^ target genes and gene expression in *fru P1*-expressing neurons (DALTON *et al*. 2013; PALMATEER *et al*. 2023). Gene ontology enrichment analyses was performed on the gene lists (ASHBURNER *et al*. 2000; LYNE *et al*. 2007; ALEKSANDER *et al*. 2023; HU *et al*. 2023). The enrichment analyses, gene lists, and statistical tests are provided (**Supplementary Table 4**)

### Olfactory/Gustatory Receptor-expressing neuronal silencing screen

We obtained *Gal4* driver lines that have been shown to be expressed in *Or/Gr*-expressing neurons (DUNIPACE *et al*. 2001; COUTO *et al*. 2005; FISHILEVICH *et al*. 2005; FISHILEVICH AND VOSSHALL 2005; KWON *et al*. 2011), for 13/15 genes that were identified as *D. melanogaster*-specific from the CUT&Tag analyses (**see strain list: Supplementary Table 1**). Males from these *Gal4* lines were each crossed to virgins of the genotype *UAS<stop<Kir2.1; fru^Flp^*, to generate males with the respective *Or/Gr* ∩ *fru P1*-expressing neurons silenced. Control lines were made by crossing male Canton S flies to virgin *UAS<stop<Kir2.1; fru^Flp^*, and by crossing Canton S virgin flies to each respective *Or/Gr*-*Gal4* line. We performed a choice assay, with males from these crosses in a chamber with two female virgins: one *D. simulans* (red eyes) and one *D. melanogaster* (white eyes). Based on two independent people watching the blinded videos for the *Or/Gr-*Gal4 ∩ *fru P1*< *<Kir2.1* genotypes, seven of the genotypes with silenced *Or/Gr* ∩ *fru P1*-expressing neurons were identified as potentially having changes in the amount of time courting *D. simulans* females, compared to controls. The videos for these seven genotypes and controls were coded using the Noldus Observer XT and the following behaviors were coded: following, wing extension, switching, copulation attempt, and successful copulation. Each video was coded twice: once coding for any behavior performed towards the *D. melanogaster* (white eyed) and once coding for any behavior performed towards the *D. simulans* (red eyed). An n of ∼30 videos were coded blind.

## Results

### Courtship behaviors of *D. melanogaster* and *D. simulans*

To evaluate hybrid courtship behaviors in the context of their pure species parents, we examined courtship behavior of wild type *D. melanogaster* (Berlin strain), wild type *D. simulans* (C167.4 strain) toward *D. melanogaster* and *D. simulans* females (**Figure 2A-H**). Consistent with previous knowledge on species partner preferences (MANNING 1959), results from the single-pair courtship assay show that male *D. melanogaster* males actively engages in courtship behaviors toward both conspecific and heterospecific *D. simulans* females (**Figure 2A-H**). *D. melanogaster* males actively engage in courtship behaviors toward both conspecific and heterospecific (*D. simulans*) females (**Figure 2A-H**). No significant differences were detected in the median courtship index (CI), wing extension index (WEI), and number of copulation attempts. In contrast, the *D. simulans* males only courted conspecific females, with the exception of one outlier (**Figure 2A-H**). However, this outlier male was not actively engaged in any scored courtship element beyond following the female (**Figure 2A-H**).

**Figure 2.**
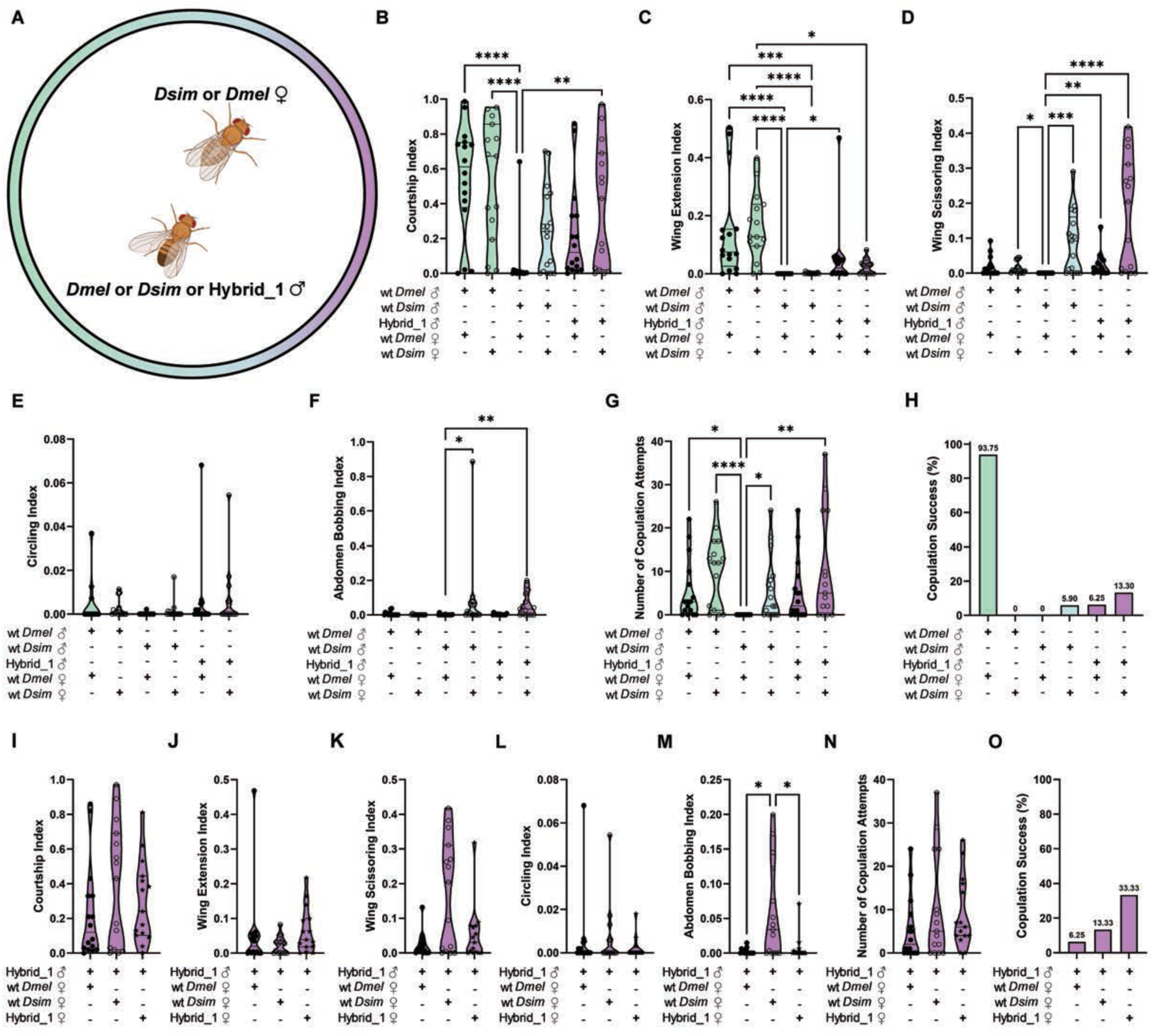
Courtship behavior of *D. melanogaster*, *D. simulans* and Hybrid_1 males. (**A**) Schematic of single-pair assay. *Created in BioRender. Arbeitman, M. (2025)* https://BioRender.com/v3me7zc. (**B-F**) Behavioral indices for pure species and Hybrid_1 males courting. The *D. melanogaster* strain is Berlin, and the *D. simulans* strain is DsimC167.4. (**B**) Courtship Index (CI), (**C**) Wing Extension Index (WEI), (**D**) Wing Scissoring Index (WSI), (**E**) Circling Index, and (**F**) Abdomen Bobbing Index (ABI). (**G**) Number of copulation attempts toward each female species. (**H**) Percentage of males that successfully copulated with each female species. (**I–M**) Behavioral indices for Hybrid_1 males courting *D. melanogaster*, *D. simulans*, and Hybrid_1 females. (**I**) CI, (**J**) WEI, (**K**) WSI, (**L**) Circling Index, and (**M**) ABI. (**N**) Number of copulation attempts toward each female target. (**O**) Percentage of Hybrid_1 males that successfully copulated. (**B–O**) Graph legends indicate pairings (“+” = included, “–” = excluded). Male species color-coded: *D. melanogaster* (green), *D. simulans* (blue), Hybrid_1 (purple). Female targets: filled points = *D. melanogaster*, open = *D. simulans*, stars = Hybrid_1. Violin plots (**B–G, I–N**) show medians and IQRs. Assays lasted 20 minutes. Statistical tests: Kruskal–Wallis ANOVA with Dunn’s post hoc. Only significant comparisons shown. Significance: *p < 0.05, **p < 0.01, ***p < 0.001, ****p < 0.0001. See Supplementary Table 2 for details.

Overall, males primarily perform species-typical behaviors. *D. melanogaster* males exhibited a significantly higher WEI toward both species of female compared to *D. simulans* males (**Figure 2C**). Although not statistically significant, *D. simulans* males showed a higher mean wing scissoring index (WSI) towards conspecific females than *D. melanogaster* males towards either of the female species (**Figure 2D**). *D. melanogaster* males had high copulation success with conspecific females (93.75%) and were unsuccessful with *D. simulans* females (**Figure 2H**). Conversely, *D. simulans* were rarely successful with conspecific females (5.90%) and had no successful copulations with *D. melanogaster* females (**Figure 2H**). Altogether, these behavioral patterns are in agreement with previously described behaviors for these species (SPIETH 1952).

We additionally tested males of two *D. melanogaster* strains (Berlin and Canton S), two *D simulans* strains (C167.4 and *Lhr*), to assess the impact of strain variation on species-specific behaviors (**Supplementary Figure 1**). Females of two *D. melanogaster* strains were also used to assess the impact of female strain background differences. There was not a significant impact of *D. melanogaster* female strain across the courtship behaviors (**Supplementary Figure 1**). Both strains of *D melanogaster* males similarly courted both *D. melanogaster* female strains, while *D. simulans* males did not court either. The two *D. simulans* male strains also similarly courted *D. simulans* females. These results indicate that across laboratory strains, pure species male courtship behaviors are robust to strain background differences in males and female targets.

### Hybrid males exhibit behaviors consistent with female target

Hybrid males (Hybrid_1; *D. melanogaster* Berlin females are parent) court both *D. melanogaster* and *D. simulans* females (**Figure 2A-H**). Consistent with previous studies, they perform all courtship behaviors of their pure species parents and attempt to copulate with either target (**Figure 2A-H**) (WOOD AND RINGO 1980; WOOD *et al*. 1980). Interestingly, Hybrid_1 males show a higher WEI toward *D. melanogaster* than towards *D. simulans* females, while exhibiting more pronounced scissoring behavior and bobbing toward *D. simulans* females. This is the most evident for the scissoring behavior (WSI), which *D. simulans* males perform vigorously in our assays (**Figure 2A-H**) (SPIETH 1952).

If we examine hybrid male behavior towards hybrid females, hybrid males court hybrid females most similarly to how they court *D. melanogaster* females (**Figure 2I-O and Supplementary Figure 1**). Abdomen bobbing and scissoring behaviors are highest towards *D. simulans* females than towards *D. melanogaster* and Hybrid_1 females. However, Hybrid_1 males show higher mean wing extension towards Hybrid_1 females than towards *D. melanogaster*. Hybrid females have a pheromone profile similar to that of *D. melanogaster*, as they also produce 7, 11-HD (SHIRANGI *et al*. 2009). This suggests that hybrid males are differentiating between the female targets based on their pheromone profile. Taken together, the potential for both *D. melanogaster* and *D. simulans* behaviors are built into the hybrid male nervous system, and differences in the female target elicit distinct behavioral outputs.

### Hybrid male courtship preference and behavioral plasticity

In the single-pair courtship assays, Hybrid_1 males showed higher mean CI, WSI, ABI, and circling index toward *D. simulans* females (**Figure 2**). Given these observations, we next investigated whether hybrid males show a preference when presented with both female species in a courtship preference assay (**Figure 3A-C**), with the female species distinguished by eye color (red or white). Both *D. melanogaster* males and *D. simulans* males exhibited a significant courtship, wing extension, and wing scissoring preference toward conspecific females with wild type red eyes (**Figure 3D-G**). However, when the *D. melanogaster* female has white eyes, *D. melanogaster* males show no significant preference for *D. melanogaster* females across courtship, wing extension, and scissoring, suggesting that the presence of *D. melanogaster* pheromones is not sufficient to ensure species recognition when the female adult phenotypes are altered. In contrast, *D. simulans* males still prefer conspecific females, even if they have white eyes, consistent with pheromones being a critical distinguishing factor for *D. simulans* males.

**Figure 3.**
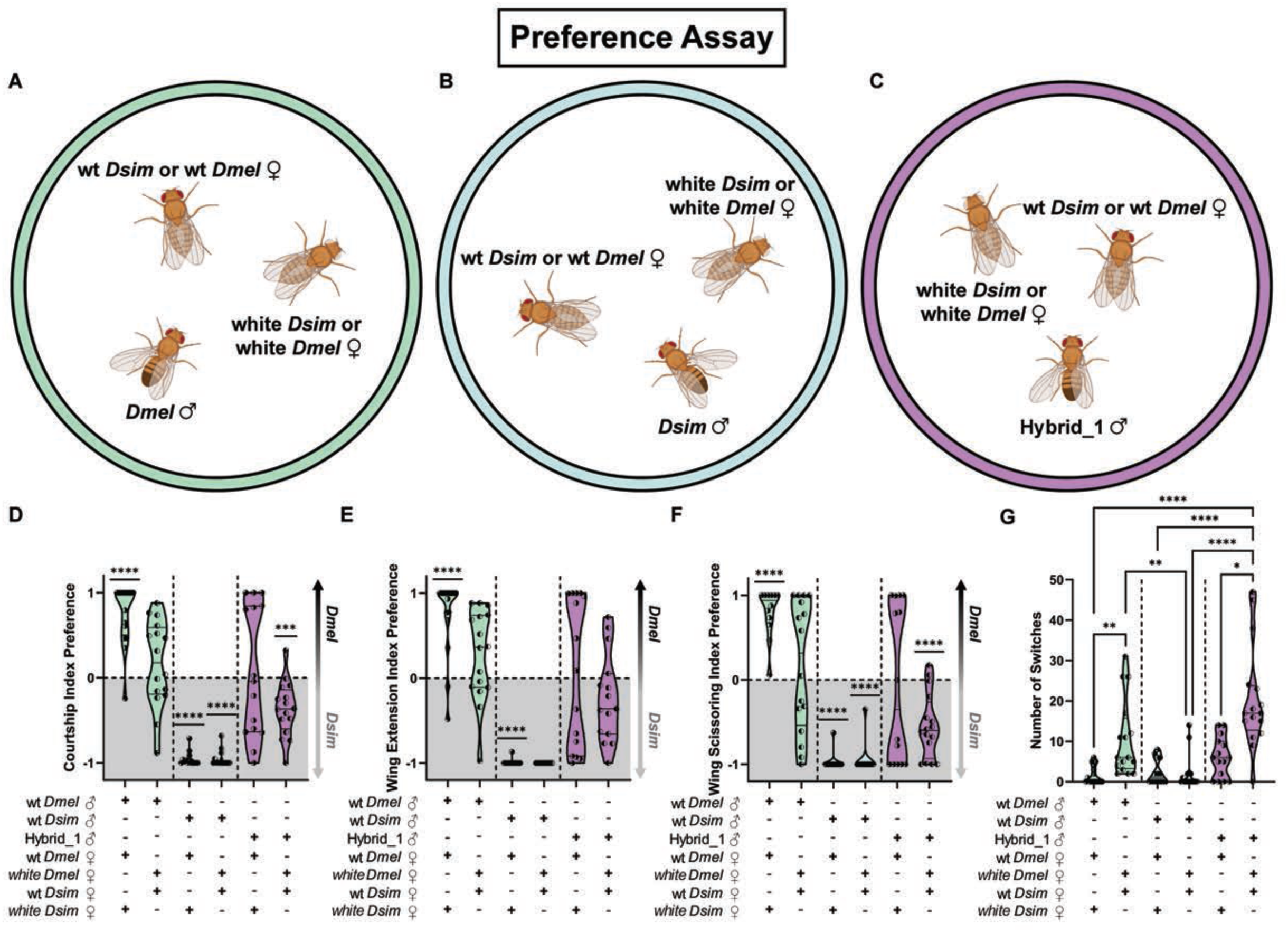
Courtship behavior preference assays of *D. melanogaster*, *D. simulans*, and and Hybrid_1 males. (**A–C**) Diagram of preference assay with one male and two female virgin targets. Male strains: *D. melanogaster* (Berlin), *D. simulans* (DsimC167.4). *Created in BioRender. Arbeitman, M. (2025)* https://BioRender.com/v3me7zc. (**D–F**) Behavioral preference indices for males toward *D. melanogaster* (Berlin or white eyed w;Berlin) and *D. simulans* (DsimC167.4 or white eyed Simw105) females. (**D**) Courtship Index Preference. (**E**) Wing Extension Index Preference. (**F**) Wing Scissoring Index Preference. Scores >0 indicate preference for *D. melanogaster*, <0 for *D. simulans*. (**G**) Number of courtship switches between female targets. (**D–G**) Pairings indicated below each graph (“+” = included, “–” = excluded). Male genotypes color-coded: *D. melanogaster* (green), *D. simulans* (blue), Hybrid_1 (purple). Female targets shown with half-filled symbols: open-left = wild-type *D. melanogaster* + white *D. simulans*, open-right = white *D. melanogaster* + wild-type *D. simulans*. Violin plots show medians and IQRs. Assays lasted 20 minutes. Statistical tests: (D–F) one-sample t-tests; (G) Kruskal–Wallis ANOVA with Dunn’s post hoc. Only significant comparisons shown. Significance: *p < 0.05, **p < 0.01, ***p < 0.001, ****p < 0.0001. See Supplementary Table 2 for details.

Hybrid_1 males do not show a significant courtship or wing extension preference toward either pure-species female, when *D. simulans* females have white eyes (**Figure 3D-G**). However, Hybrid_1 males did show a significant courtship and wing scissoring preference toward *D. simulans* females when *D. simulans* females have red eyes. Previous studies have shown an impact of eye color on courtship in *D. melanogaster* (MCKINNEY AND BEN-SHAHAR 2019).

We also observed the highest frequency of switching between the courtship targets in Hybrid_1 males when the preference assay was performed with white eyed *D. melanogaster* females (**Figure 3G**). Visual inspection of the courtship preference videos showed that the Hybrid_1 males would engage in courting the *D. melanogaster* female with *D. melanogaster*-like behaviors, performing unilateral wing extensions toward her. In the same assay, when proximal to the *D. simulans* female, the hybrid male would rapidly switch his behavior toward her to more *D.simulans*-like behaviors, such as circling the female, abdomen bobbing and scissoring his wings (**Video 1**). The data show that there is female target-dependent plasticity in the hybrids, with the potential for both species behaviors built into the nervous system. This suggests that in hybrids the courtship sequence is not generated as fixed, sequence-dependent action pattern but can be behaviorally tuned based on the female target. The data also suggests that there are multiple cues directing courtship preference that are integrated to elicit different behavioral outcomes, for both the pure species and hybrid males, with eye color and pheromones making contributions.

### Hybrid strain genetic background impacts male courtship behavior

Next, we examined the impact of the strain background of the *D. melanogaster* female parent of the hybrid male on hybrid male behavior. We generated hybrid males with females that readily mate with *D. simulans Lhr* males. Among several wild type *D. melanogaster* strains tested, we found a strain originally derived from *white^1118^*, but with *w^+^* introgressed back into the strain background, that readily mated with *D. simulans Lhr* males. The hybrid males generated from this cross are called Hybrid_2 and were assayed in both single-pair and preference assays, as above (**Figure 4A-C**). The behaviors of Hybrid_2 differed from those of Hybrid_1, exhibiting substantially less overall courtship towards *D. melanogaster* females (**Figure 4D-H**). Similar to Hybrid_1, female eye color impacted female species preference, with Hybrid_2 males showing significantly more courtship and a preference towards *D. simulans* females, but primarily when *D. simulans* females had red eyes (**Figure 4I-Q**). A similar impact of strain on hybrid behavior has been observed (SHAHANDEH *et al*. 2020). The results show that the hybrids are also plastic in their behavioral outcomes, based on the strain background of the parents, in a way that is not observed in pure species. This can be understood within the framework of considering sex-specific phenotypes as quantitative traits, with differences in the genetic backgrounds having a larger influence in a hybrid strain genetic environment, than in pure species backgrounds where courtship behavior is more robustly specified.

**Figure 4.**
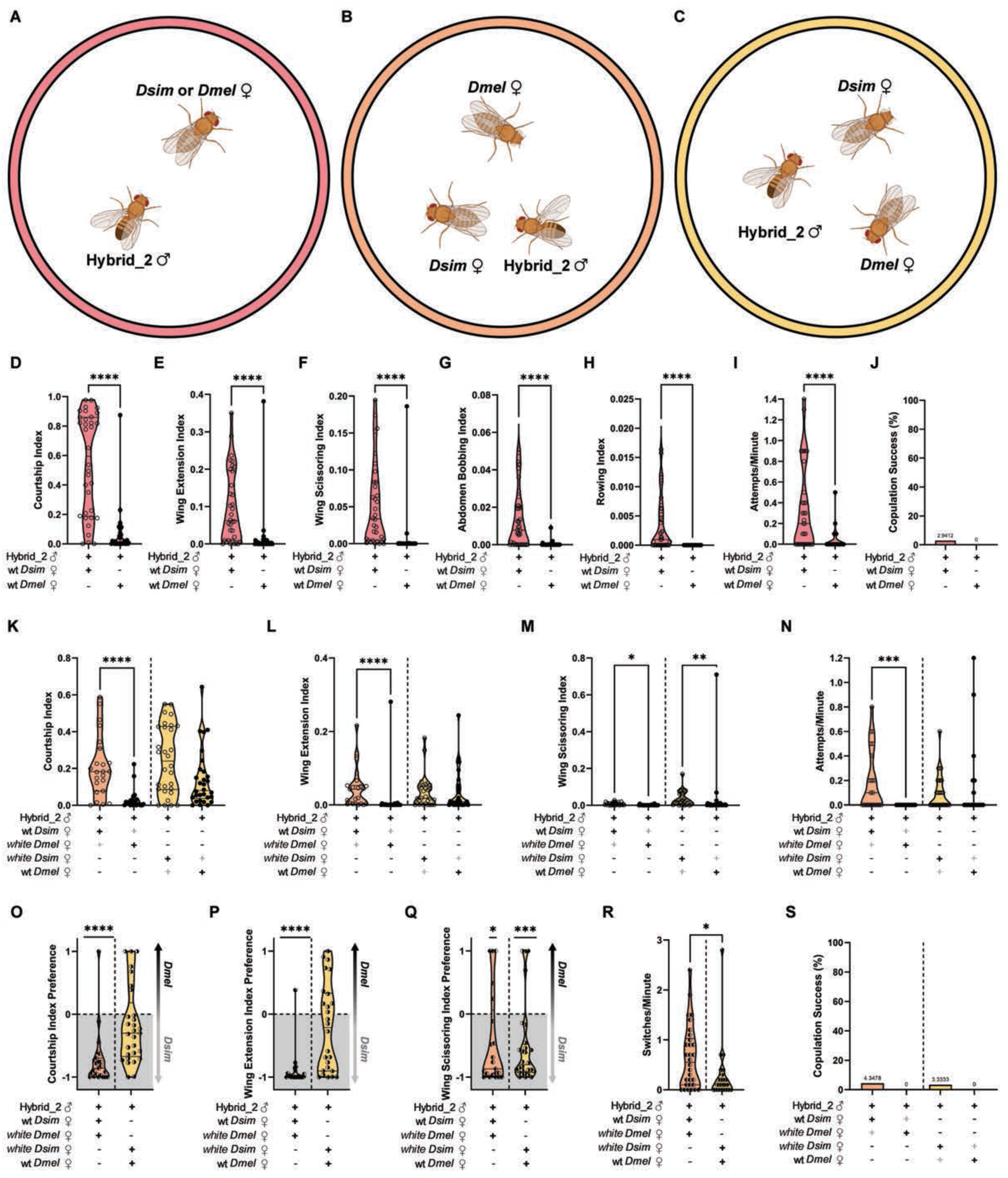
Courtship behavior for Hybrid_2 males in single-pair assays and preference assays. (**A**) Schematic of single-pair assay. *Created in BioRender. Arbeitman, M. (2025)* https://BioRender.com/v3me7zc. (**B-C**) Diagram of preference assay, with one male and two virgin females. (**D-H**) Male behavior indices for Hybrid_2 males toward *D. melanogaster* (*w*+WG) or *D. simulans* (DsimC167.4) females. (**D**) Courtship Index (CI), (**E**) Wing Extension Index (WEI), (**F**) Wing Scissoring Index (WSI), (**G**) Abdomen Bobbing Index (ABI), and (**H**) Rowing Index (RI). (**I**) The rate at which the male attempted copulation toward each species of female per minute. (**J**) Percentage of males that successfully copulated with females of each species. (**K-M**) Male behavior preference assay indices toward *D. melanogaster* (*w*+WG or *w*; Canton S) females or *D. simulans* (DsimC167.4 or Simw^195^) females. (**K**) Courtship Index (CI), (**L**) Wing Extension Index (WEI), and (**M**) Wing Scissoring Index (WSI). (**N**) The rate at which the male attempted copulation toward each species of female per minute. (**O-Q**) Male behavior preference assay indices toward *D. melanogaster* (w+WG or w; Canton S) females and *D. simulans* (DsimC167.4 or Simw^195^) females for (**O**) Courtship Index Preference, (**P**) Wing Extension Index Preference, and (**Q**) Wing Scissoring Index Preference. A score greater than 0 indicates a preference towards *D. melanogaster* females, while a score less than 0 indicates a preference towards *D. simulans* females. (**R**) The rate at which Hybrid_2 males switched between courting *D. melanogaster* and *D. simulans* females during preference assay per minute. (**S**) Percentage of males that successfully copulated with females of each species during preference assay. (**D-S**) Graph legends indicate male and female pairings (“+” = included, “–” = not included; light gray “+” = present but data not shown). Color coding: pink = single-pair assays; orange = preference assays with wild-type *D. simulans* and white *D. melanogaster*; yellow = wild-type *D. melanogaster* and white *D. simulans*. Female species are symbolized by point fill: filled = *D. melanogaster*, open = *D. simulans*, half-filled left = wild-type *D. melanogaster* + white *D. simulans*, half-filled right = white *D. melanogaster* + wild-type *D. simulans*. (**D–I, K–R**) Violin plots show median and IQR. (D–I, R) Mann-Whitney tests; (K–N) Kruskal-Wallis with Dunn’s post hoc; (**O–Q**) one-sample t-tests. Only significant comparisons shown. Significance: *p < 0.05, **p < 0.01, ***p < 0.001, ****p < 0.0001. See Supplementary Table 2 for details.

### Examination of subpopulations of *fru P1*-expressing neurons in *D. melanogaster* and hybrid males

The *fru P1* neuronal circuitry that underlies hybrid courtship behavior has not been examined, leaving open the question as to how hybrid males can perform a novel combination of behavioral repertoires. One possibility is that hybrid behaviors arise from small-scale quantitative differences in the *fru P1* neuroanatomical substrate, rather than large scale changes in neuroanatomy, as is observed for comparisons between the species (SEEHOLZER *et al*. 2018) (reviewed in ANHOLT *et al*. 2020). To gain insight into this, we examine Fru^M^ expression, and *fru P1*-expressing neurons in hybrid males. First, we compared *fru P1* and Fru^M^ expression between hybrid and *D. melanogaster* males (**Figure 5A-B**). We used *fru P1*-*Gal4* (MANOLI *et al*. 2005) to drive expression of *UAS-mCherry.NLS* and stained with anti-Fru^M^ antibody in both *D. melanogaster* and hybrid males. This revealed highly congruent expression patterns (**Figure 5A-B**), consistent with *fru P1*-expression being similar between the species (SEEHOLZER *et al*. 2018).

**Figure 5.**
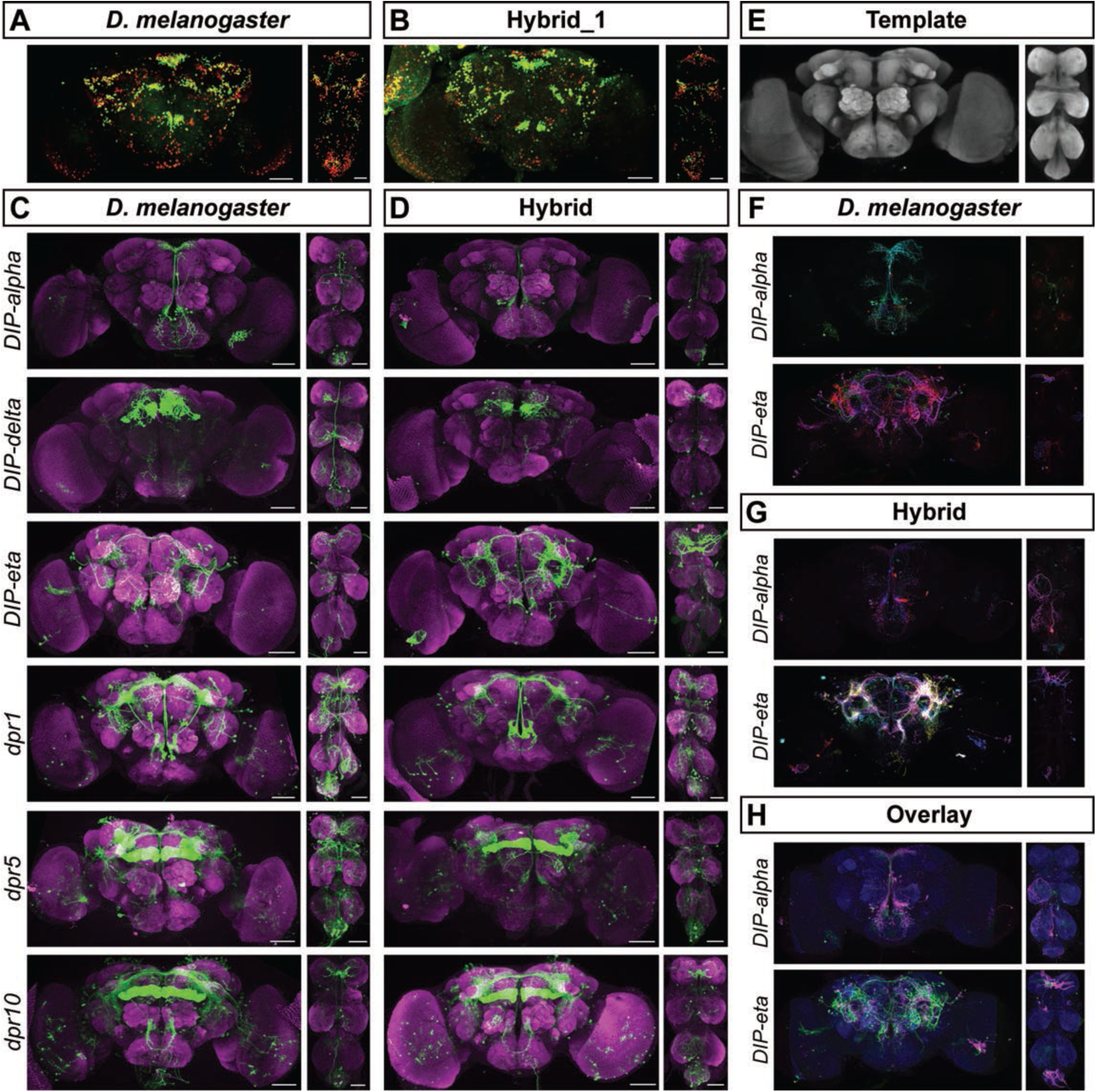
*fru P1 ∩ dpr/DIP*-expressing neurons are similar in *D. melanogaster* and hybrid males. (**A–B**) *fru P1-Gal4 > UAS-mCherry.NLS* (red) and Fru^M^ antibody (green) expression in brain (left) and VNC (right) of (**A**) D. melanogaster and (**B**) hybrid males. (**C–D**) *dpr/DIP ∩ fru P1* neurons in brain and VNC of (**C**) D. melanogaster and (**D**) hybrid males. *Gal4* drivers indicated at left. DIP-alpha, DIP-delta, dpr5, and dpr10 samples from 4–7-day old adults; DIP-eta and dpr1 from 0–24 hr old adults. Scale bars = 50 μm. (**E**) *D. melanogaster* and hybrid male template brain and VNC. *D. melanogaster* and hybrid average brain and VNC generated using anti-Nc82 antibody staining for non-rigid image registration. (**F-H**) *D. melanogaster* and hybrid male *DIP-alpha ∩ fru P1* and *DIP-eta ∩ fru P1* neurons visualized after registration. (**F**) On top, *DIP-alpha ∩ fru P1* expression in *D. melanogaster* males from multiple individuals, brain n = 5 and VNC n = 3. On bottom, *DIP-eta ∩ fru P1* expression in *D. melanogaster* males from multiple individuals, brain n = 4 and VNC n = 2. (**G**) On top, *DIP-alpha ∩ fru P1* expression in hybrid males from multiple individuals, brain n = 4 and VNC n = 3. On bottom, *DIP-eta ∩ fru P1* expression in hybrid males from multiple individuals, brain n = 7 and VNC n = 2. (**F-G**) Different colors show individual expression patterns. (**H**) On top, *D. melanogaster* (green) and hybrid (magenta) *DIP-alpha ∩ fru P1* patterns overlaid on template brain and VNC. n = 3 individual expression patterns from *D. melanogaster* and hybrids males for brain and VNC. On bottom, *D. melanogaster* (green) and hybrid (magenta) *DIP-eta ∩ fru P1* patterns overlaid on template brain and VNC. n = 3 individual expression patterns from *D. melanogaster* and hybrids males for brain and n = 2 for VNC.

To examine potential anatomical differences with higher resolution, we examine smaller populations of neurons that we previously showed are important for producing male courtship behavior in *D. melanogaster* (BROVERO *et al*. 2021). We leverage tools we built in our recent study to examine the role of *fru P1* neurons that co-express genes encoding immunoglobulin domain super family (IgSF) proteins — *defective proboscis extension response* (*dpr)* and their interacting partners *dpr interacting proteins* (*DIPs*). To subdivide *fru*-expressing neurons using *dpr*/*DIP* expression, we used a genetic intersectional strategy. This approach combines the *GAL4*/*UAS* binary system with the *Flp*/*FRT* recombinase system to express *mCD8::GFP* only in *dpr*/*DIP* and *fru P1* co-expressing neurons (*dpr*/*DIP* ∩ *fru P1* neurons). Here, we evaluate co-expressing patterns for those with spatially restricted subsets (*DIP-alpha* ∩ *fru P1* and *DIP-delta* ∩ *fru P1*), as well as those with broader co-expression patterns (*dpr1* ∩ *fru P1*, *dpr10* ∩ *fru P1*, and *DIP-eta* ∩ *fru P1*) (BROVERO *et al*. 2021). When we compare these *dpr*/*DIP* ∩ *fru P1* neurons between *D. melanogaster* and hybrid males, the morphology appears conserved, though there is individual variation in both *D. melanogaster* and hybrids (**Figure 5C-D**).

To facilitate direct comparisons, we aligned the expression patterns from both *D. melanogaster* and hybrid males into a standardized image space using non-rigid image registration (ROHLFING AND MAURER 2003; ROHLFING 2012). We directly compare *DIP-alpha* ∩ *fru P1* expression, which is spatially restricted, and *DIP-eta* ∩ *fru P1* expression, which has a broader expression pattern. For image registration, we generated an average brain and ventral nerve cord (VNC) using images from both *D. melanogaster* and hybrid males as a registration template (**Figure 5E**). We registered the highest quality images, resulting in a smaller number of images used for this analysis. The registration shows the expression stochasticity between flies, with neurons from multiple individuals shown in different colors for both *D. melanogaster* and hybrids (**Figure F-H**). Overlaying the *D. melanogaster* expression patterns (green) with the hybrid expression patterns (magenta) revealed largely congruent overall expression (**Figure F-H**), with similar levels of variation observed within the data from *D. melanogaster* or hybrids.

### Identification of Fru^M^ target genes in *D. melanogaster* and *D. simulans*

The behavioral, anatomical, and physiological data from the pure species and hybrids suggests that differences in Fru^M^ target genes, within a largely shared neuroanatomical *fru P1*-expressing neuroanatomical substrate, could contribute to explaining species differences in behaviors. This could also explain how hybrids have the potential for both species behaviors built into their nervous system, as both species target genes are present in the hybrids. Moreover, the idea that species differences may be driven by differences in Fru^M^ target genes is consistent with a previous study that showed that species differences in behavior does not result from evolution of the *fru* and *dsx* loci (CANDE *et al*. 2014).

To investigate the molecular basis of species-specific behaviors, we identified Fru^M^ target genes in *D. melanogaster* and *D. simulans* using the CUT&Tag approach, modified for whole brain tissues (KAYA-OKUR *et al*. 2019; AHMAD 2020). We identified bound peak regions using the SEACR algorithm (MEERS *et al*. 2019), mapping the Illumina sequencing reads from both species to the *D. melanogaster* genome. This genome was selected due to its high sequence conservation with *D. simulans* and superior annotation compared to *D. simulans* (CLARK *et al*. 2007; OZTURK-COLAK *et al*. 2024). Although this approach may miss *D. simulans*-specific genes, this is balanced and offset by the ability to provide annotation for peaks identified in both species. Two independent CUT&Tag libraries were generated for each species, with negative controls to facilitate peak calling. A gene was considered a Fru^M^ target if identified in both replicates for the species. This resulted in 4,415 and 5,097 Fru^M^ target genes in *D. melanogaster* and *D. simulans*, respectively (**Figure 6A and Supplementary Table 3**). Of these, 2,875 genes were common to both species, while 1,541 and 2,223 genes are potentially species-specific for *D. melanogaster* and *D. simulans*, respectively (**Figure 6A**).

**Figure 6.**
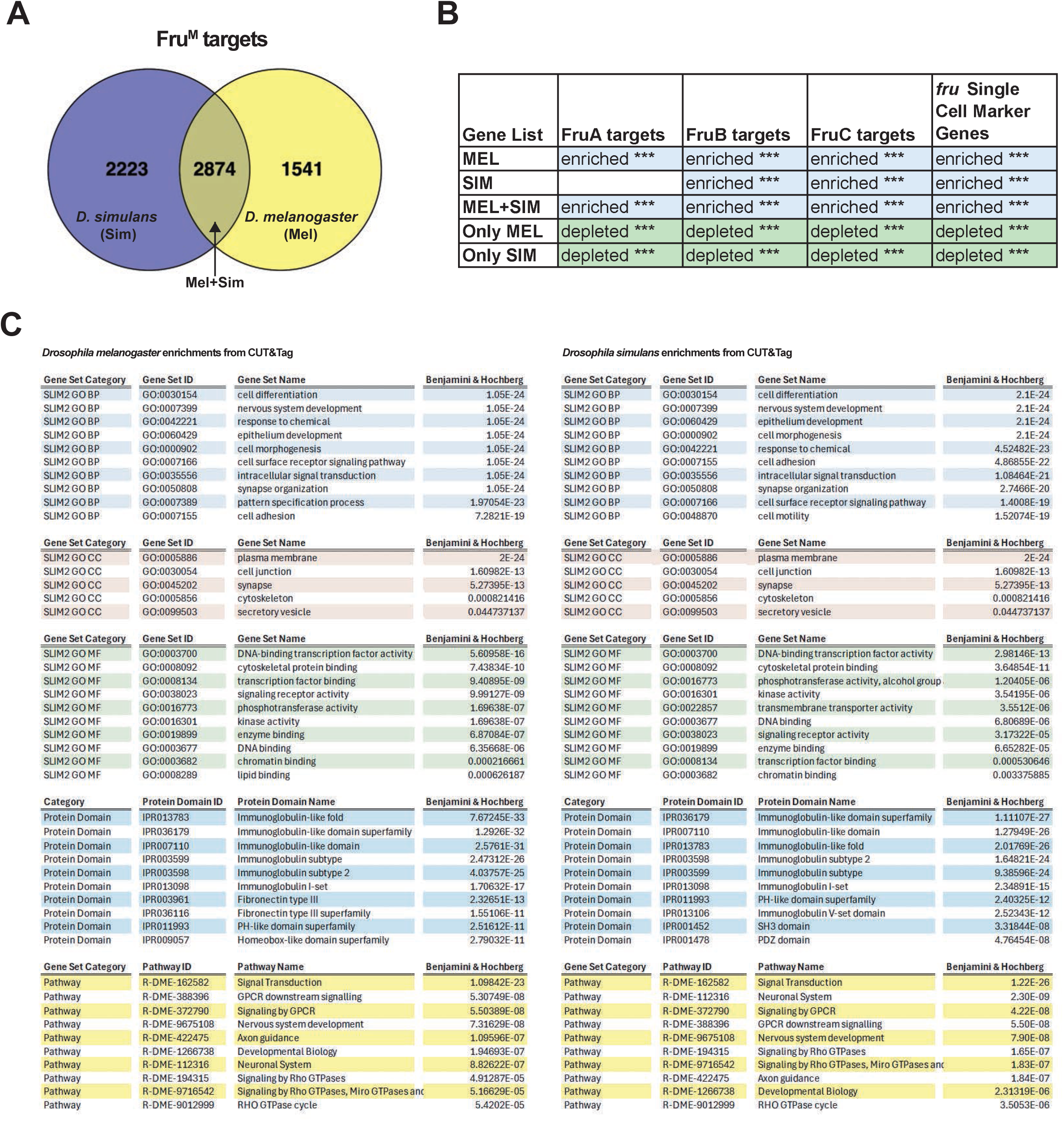
Analyses of *D. melanogaster* and *D. simulans* Fru^M^ target genes identified using the CUT&Tag approach. (**A**) Venn diagram showing overlap of Fru^M^ target genes identified in *D. melanogaster* (Mel), *D. simulans* (Sim), and shared targets (Mel+Sim) (see Supplementary Table 3). (**B**) Overlap of CUT&Tag genes lists (**A**), with genes with Fru^M^ binding sites (DALTON *et al*. 2013), and marker genes from a single-cell RNA sequencing study of *fru-P1*-expressing neurons (PALMATEER *et al*. 2023) (**Supplementary Table 4**). (**C**) Gene Ontology analyses of gene lists from (**A**).

To validate the CUT&Tag approach and gain further insight about the identified Fru^M^ target genes, we compared the gene lists to other genomic data sets (**Figure 6B**). We first examined if the Fru^M^ target genes are enriched with those that we previously identified as having a Fru^M^ binding site. The *fru P1* transcripts (*P1* promoter) that produce Fru^M^ isoforms are sex-specifically spliced at the 5’ end to generate the Fru^M^ isoforms. Alternative splicing at the 3’ end of all *fru* transcript classes results in inclusion of different alternate DNA-binding domain encoding exons, with the A, B, and C DNA binding domains being the predominant ones in the nervous system. Our prior SELEX experiments identified the distinct DNA binding motifs for these domains (DALTON *et al*. 2013).

We examined overlaps between Fru^M^ target genes from CUT&Tag in *D. melanogaster*, *D. simulans*, and both species, with the gene lists for those that have Fru A, B, and C DNA binding motifs (**Figure 6B**). We find that the Fru^M^ target genes are significantly enriched with genes having the Fru^M^ binding motifs. The only exception being the genes with the Fru^A^ motif and the Fru^M^ target genes identified in *D. simulans*, though there is not a significant depletion, as is observed in the identified species-specific targets (see below). This supports the idea that the Fru^M^ target genes identified with the CUT&Tag approach are valid. In addition, given that genes with the A, B, and C DNA binding motif are all enriched and/or found at high numbers in the Fru^M^ target genes found in *D. melanogaster*, and *D. simulans* suggests that the Fru DNA binding domains have not evolved to have highly different functions between the species; this conservation is predicted based on previous work (PARKER *et al*. 2014; BAKER *et al*. 2024). In contrast, species-specific Fru^M^ target genes are significantly depleted from the gene lists for those that have Fru A, B, and C DNA binding motifs (**Figure 6B**). If the species-specific Fru^M^ target genes are bona fide targets, this is potentially through binding different Fru DNA binding site motifs than those identified in our SELEX experiments (DALTON *et al*. 2013), or by binding to DNA indirectly, potentially with species-specific heterodimer partners.

We also observed significant overlap of the of Fru^M^ target genes found in *D. melanogaster*, *D. simulans* and those in common between the species, with the genes identified as “marker genes” in our single cell RNA-seq analyses of *fru P1* neurons (PALMATEER *et al*. 2023). Marker genes are defined as those having higher expression in a set of *fru P1* neurons, compared to the full set of *fru P1* neurons in the single cell RNA-seq analyses. The marker genes also molecularly define differences across subsets of *fru P1* neurons in the data set. However, the species-specific Fru^M^ target genes do not show significant overlap with the marker genes from the single-cell analyses. If the species-specific Fru^M^ targets genes are true targets, their expression profiles in the *Drosophila melanogaster* single cell RNA-seq data set are not highly specific to a sub-population of *fru P1* neurons, leaving their functional roles unclear.

Gene Ontology (GO) analyses of Fru^M^ target genes in *D. melanogaster* and *D. simulans* revealed shared categories, consistent with previous studies on *fru P1* neurons and Fru^M^-regulated expression (GOLDMAN AND ARBEITMAN 2007; DALTON *et al*. 2009; LEBO *et al*. 2009; DALTON *et al*. 2013; NEVILLE *et al*. 2014; VERNES 2014; BROVERO *et al*. 2021; BROVKINA *et al*. 2021; PALMATEER *et al*. 2021). For example, we see the “Immunoglobulin-like domain” is an enriched protein domain, a family we have shown has a role in *fru*-expressing neurons (GOLDMAN AND ARBEITMAN 2007; BROVERO *et al*. 2021). The GO analyses on the species-specific gene lists did not identify significant terms under our statistical criteria, so we are not able to ascertain what processes they may underlie. Given the role of pheromones and their receptors in species recognition, we examined Fru^M^ target gene lists for differences in genes annotated in FlyBase as “Chemosensory” and “Ion Channels” (OZTURK-COLAK *et al*. 2024)ß, with the goal of undertaking a genetic screening approach to find additional genes involved in species recognition. These are also reported as a resource for future work (**Supplementary Table 4**). We chose to focus on the 15 chemosensory genes that were identified as *D. melanogaster* specific Fru^M^ targets, which include mainly *odorant receptor* (*Or*) and *gustatory receptor* (*Gr*) genes, some of which have been previously implicated in impacting courtship based on the olfactory/taste responses (AUER AND BENTON 2016). There are well-characterized *Gal4* driver transgenic strains for 13 of these genes in *D. melanogaster*, motivating our choice of these genes for the genetic screen (DUNIPACE *et al*. 2001; COUTO *et al*. 2005; FISHILEVICH *et al*. 2005; FISHILEVICH AND VOSSHALL 2005; KWON *et al*. 2011).

### Genetic screen for role of neurons in mate discrimination that express *Or/Gr* genes identified as *D. melanogaster*-specific Fru^M^ targets

While many of the *Or* and *Gr* gene functions have been characterized in *D. melanogaster*, the roles of the 15 *Or* and Gr genes that were identified as *D. melanogaster* specific Fru^M^ targets remain unexplored in the context of species mate recognition. Given the *Or/Gr* genes are predicted to be Fru^M^ targets, we used a genetic intersectional genetic approach to either silence (*UAS<stop<Kir2.1*) or visualize (UAS<stop<visual marker gene) neurons co-expressing the *Or/Gr* and *fru P1* (*Or/Gr ∩ fru P1*) (**Figure 7A**). The behavioral screen aimed to detect differences in species preferences using a choice assay between *D. simulans* females and *D. melanogaster* females. We chose our screening conditions based on the observation that the *D. melanogaster* transgene control strain (*UAS<stop<Kir2.1/+; fru^Flp^/+*), without the *Or/Gr Gal4*, showed a significant preference for white-eyed *D. melanogaster* females, over red-eyed *D. simulans* females, for Courtship Index (CI), Wing Extension Index (WEI), and copulation attempts per minutes. We screened for changes in this preference when the *Or/Gr ∩ fru P1*-expressing neurons were silenced via expression of *Kir2.1* (reviewed in HODGE 2009). This preference assays also allowed us to determine if changes in courtship towards *D. simulans* females was a result of overall changes in courtship.

**Figure 7.**
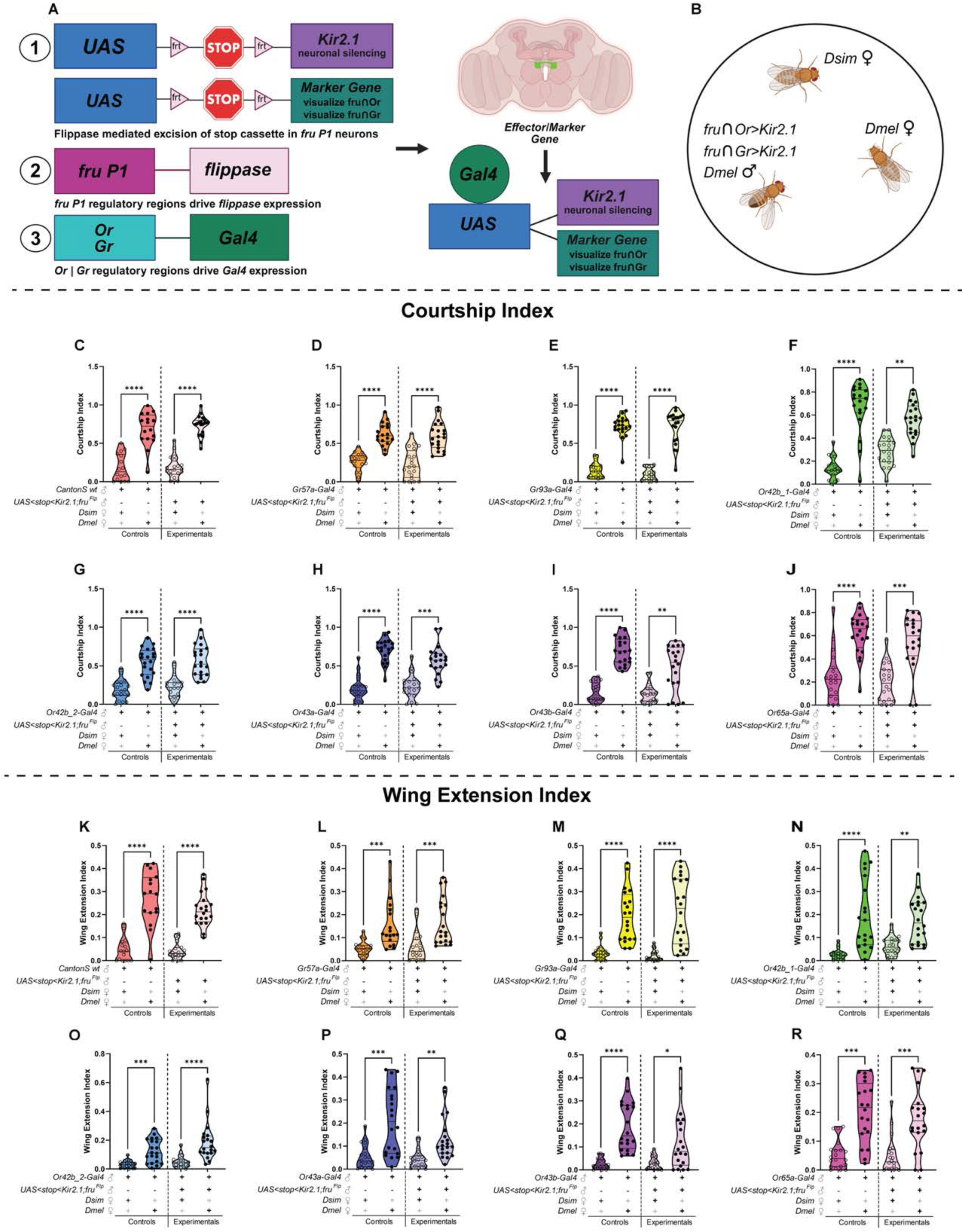
Silencing *fru P1* ∩ *odorant* and *gustatory receptor* neurons does not alter time spent courting or wing extension in a preference assay. (**A**) Genetic intersectional strategy using *fru P1*-flippase and *Or/Gr* receptor-Gal4 to express *Kir2.1* and visualize fru P1 ∩ receptor neurons. Expression requires both flp-mediated stop cassette removal and Gal4 activity. (**B**) Schematic of preference assay: control or experimental male placed with white-eyed *D. melanogaster* and red-eyed *D. simulans* females. (**C–J**) Courtship Index; (**K–R**) Wing Extension Index toward *D. melanogaster* (w; Canton S) or *D. simulans* (*DsimC167.4*) females. Graph legends indicate pairings (“+” = included, “–” = excluded); bold “+” indicates species shown. Female targets: filled points = *D. melanogaster*, open = *D. simulans*. Graph shading: darker = control, lighter = experimental. Color coding by receptor: wild type (red), Gr57a (orange), Gr93a (yellow), Or42b_1 (green; RRID_9971), Or42b_2 (light blue: RRID_9972), Or43a (dark blue), Or43b (purple), Or65a (pink). Violin plots show median and IQR. Kruskal–Wallis ANOVA with Dunn’s post hoc; only significant comparisons shown. Significance: *p < 0.05, **p < 0.01, ***p < 0.001, ****p < 0.0001. See Supplementary Table 2 for details.

We first conducted a visual screen to identify lines in which silencing *Or/Gr ∩ fru P1*-expressing neurons altered courtship towards *D. simulans* females, in the presence of *D. melanogaster* females. Seven lines were chosen for in-depth quantitative analysis (**Figure 7C-R**). Quantitative analyses of the courtship sub-behaviors revealed little impact on the preference for *D. melanogaster* females for overall courtship and wing extension. However, silencing three sets of *Or/Gr ∩ fru P1*-expressing neurons disrupted copulation preferences, increasing courtship toward *D. simulans females* (*Or 42b*, two Gal4 lines; *Or43a*; and *Or43b*) (**Figure 8**).

**Figure 8.**
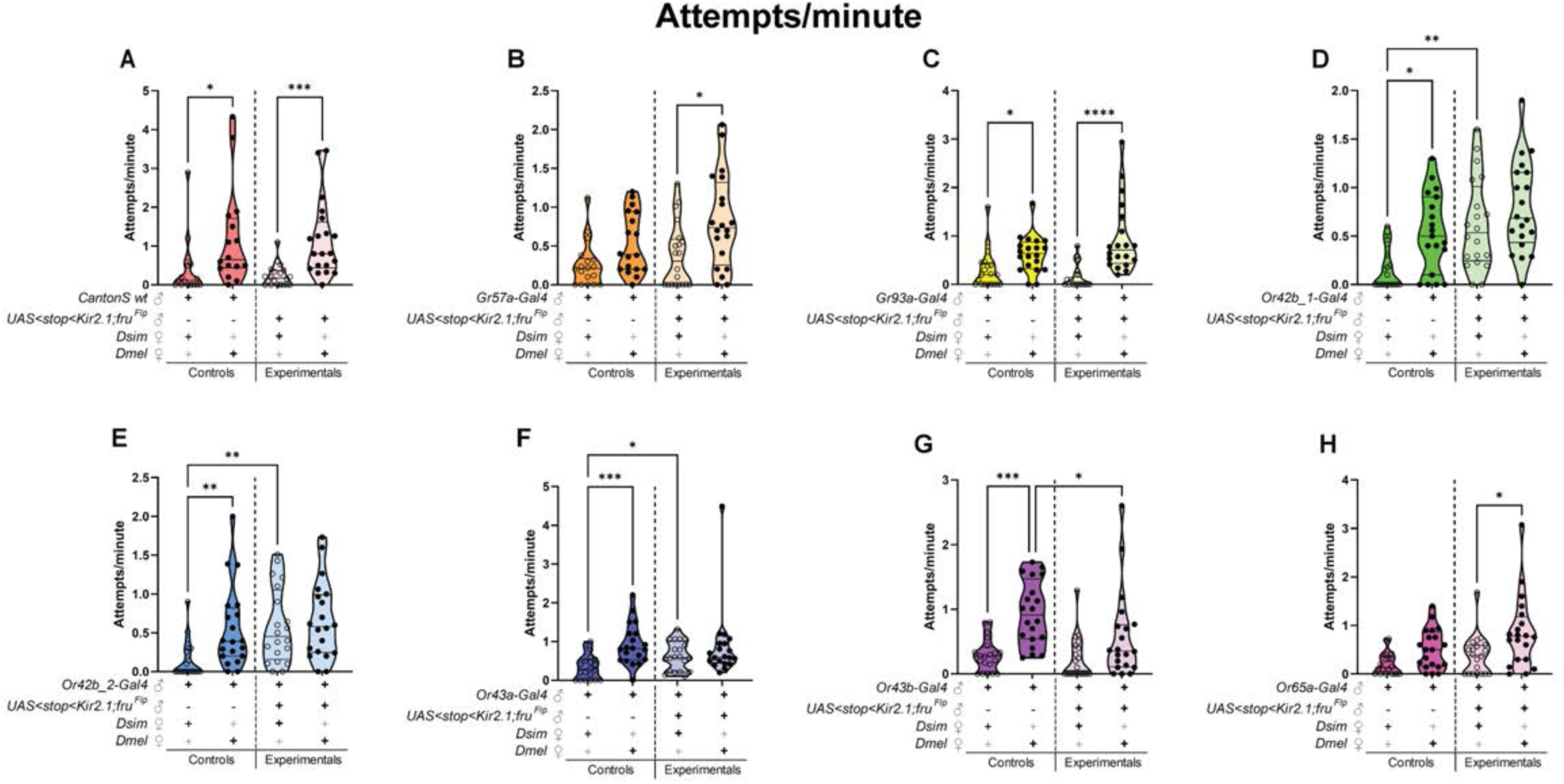
Silencing *fru P1* ∩ odorant and gustatory receptor neurons identifies Or42b, Or43a, and Or43b-expressing neurons as modulators of attempted copulation towards heterospecific females during courtship. (**A–H**) Copulation attempt rate per minute in preference assays. Pairings indicated below each graph (’+’ = included, ‘–’ = excluded; light gray ‘+’ = present but not shown). Graphs shaded by pairing: darker = wild type or Gal4 alone, lighter = Kir2.1 alone or with Gal4. Color-coded by receptor: wild type (red), Gr57a (orange), Gr93a (yellow), Or42b_1 (green), Or42b_2 (light blue), Or43a (dark blue), Or43b (purple), Or65a (pink). Female species indicated by symbols: filled = *D. melanogaster*, open = *D. simulans*. Violin plots show median and IQR. Kruskal– Wallis ANOVA with Dunn’s post hoc; only significant comparisons shown. Significance: *p < 0.05, **p < 0.01, ***p < 0.001, ****p < 0.0001. See Supplementary Table 2 for details.

Additionally, the male genotypes with *Or42b* and *Or43a ∩ fru P1*< *Kir2.1* neuronal silencing attempted to copulated more with *D. simulans* females compared to their single transgene controls (*Or-Gal4*). These olfactory receptors, which have not previously been implicated in mediating courtship behavior, appear to have a role in distinguishing conspecific from heterospecific females. Interestingly, *Or42b*-expressing neurons have previously been implicated in food preferences (SEMMELHACK AND WANG 2009; ROOT *et al*. 2011), providing a potential link to mate recognition and mating substrates.

### Co-expression of *fru P1* and *D. melanogaster*-specific Fru^M^ *Or/Gr* gene targets

To determine whether the seven *Or/Gr-Gal4* lines, selected for in-depth analyses, co-express *fru P1*, we used the genetic intersectional approach to visualize *Or/Gr ∩ fru P1*-expressing neurons (**see Figure 6A**). For these analyses we used two different Flp-out reporter strains (Multi-color: MCFO and single-color: smGDP) (**Figure 9 and Supplementary Figure 2**), which reveal different morphological features. We found that two independent *Or42b-Gal4 ∩ fru P1* genotypes target the DM1 glomeruli. *Or43a-Gal4 ∩ fru P1* targets the DA4l glomeruli, and *Or43b ∩ fru P1* targets the VM2 glomeruli. No overlap was observed with the other three Gal4 lines that did not have phenotypes in our preference assay (Or65a, Gr57a, and Gr93a).

**Figure 9.**
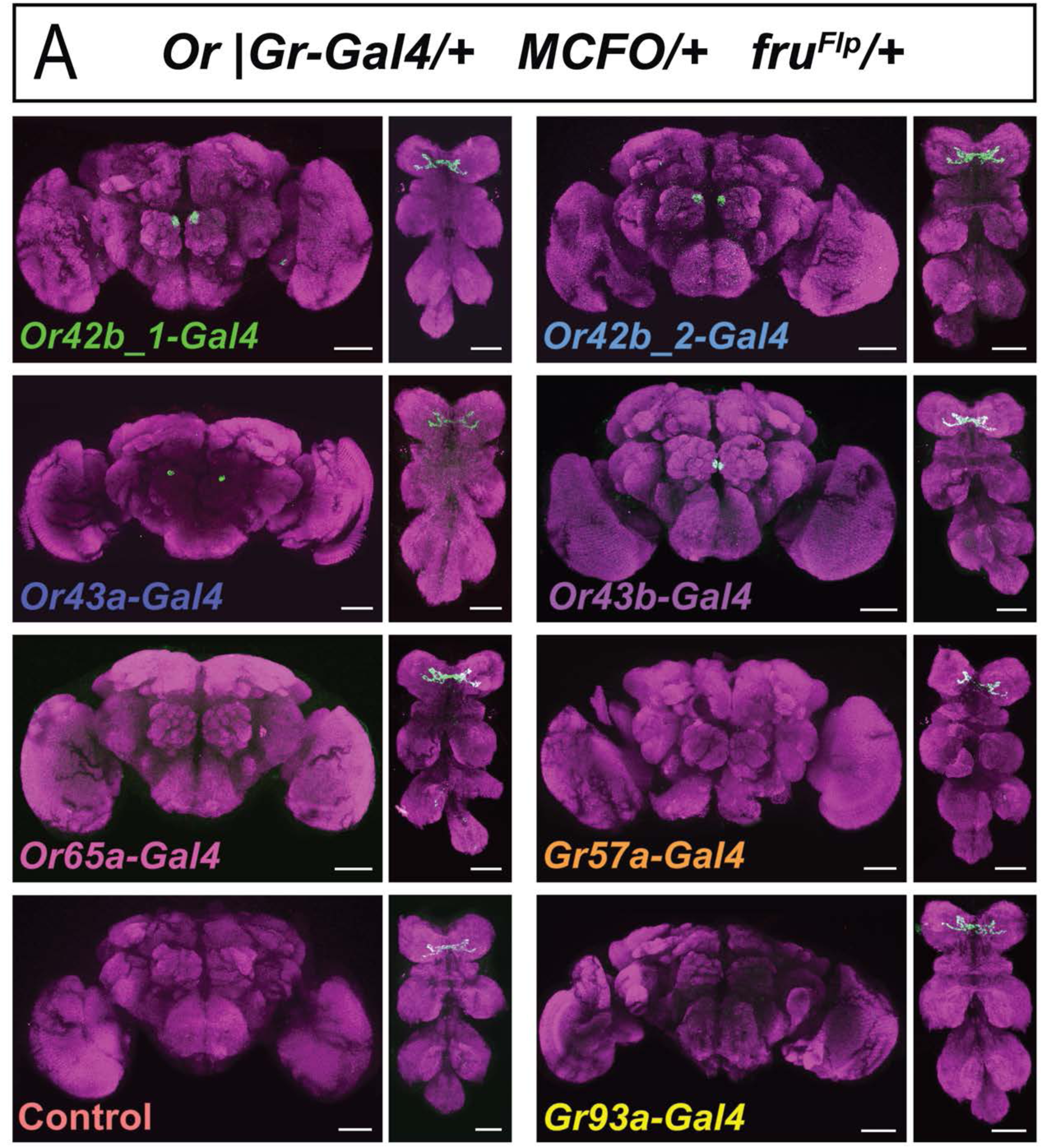
*fru P1* ∩ odorant and gustatory receptor neurons. (**A**) Confocal maximum intensity projections of brains and VNCs from 5–10-day-old adult males, using the genetic intersectional approached described in Figure 7. Genotype: *w*, *Or|Gr– Gal4/*+, *UAS<stop<MCFO*, *fru^Flp^/+.* Images show *fru P1* ∩ Or or *fru P1* ∩ Gr neurons labeled using MultiColorFlpOut (MCFO) reporters. Canton S crossed to MCFO, *fru^Flp^* is shown as control. Scale bars: 50 μm.

However, we do see clear GFP signal in the proboscis when we use Gr57a-Gal4 to drive UAS-NLS::GFP and UAS-mcd8::GFP (**Supplementary Figure 3**). The role of Or65a and Gr93 is unclear, given we did not detect staining with additional UAS-reporter genes. *fru P1* is known to be expressed in odorant receptor neurons that innervate DA1 (Or67d), Va1v (Or47b) and VL2a (Ir84a) glomeruli (MANOLI *et al*. 2005; STOCKINGER *et al*. 2005; KURTOVIC *et al*. 2007; DWECK *et al*. 2015). Additionally, low-level *fruitless* expression was detected in other classes of odorant receptor neurons during pupal stages, using single-cell RNA sequencing (MCLAUGHLIN *et al*. 2021), although it remains unclear if this is *fru P1* transcripts or the *fru* transcript classes that produce common isoforms. The results presented her reveal additional odorant receptor neurons that express *fru P1* and that have unexpected roles in species recognition and male courtship.

### Olfactory projection neurons involved in courtship innervate distinct brain regions

We used the *trans*-Tango method to visualize the post-synaptic targets of odorant receptor neurons expressing *Or42b*, *Or43a*, *Or43b* (TALAY *et al*. 2017) (**Figure 10**). Results for *Or42b* using *trans*-Tango have been previously described (TALAY *et al*. 2017). For all three odorant receptor neuron types, *trans*-Tango labeling revealed second-order local interneurons (LN) across the entire antennal lobe, with the brightest staining near the glomeruli targeted by the corresponding olfactory receptor neurons. We characterize the projection neurons using our confocal images and also cross-referenced with the FlyWire electron micrograph (EM) data (BATES *et al*. 2020), accessed through Codex (MATSLIAH).

**Figure 10.**
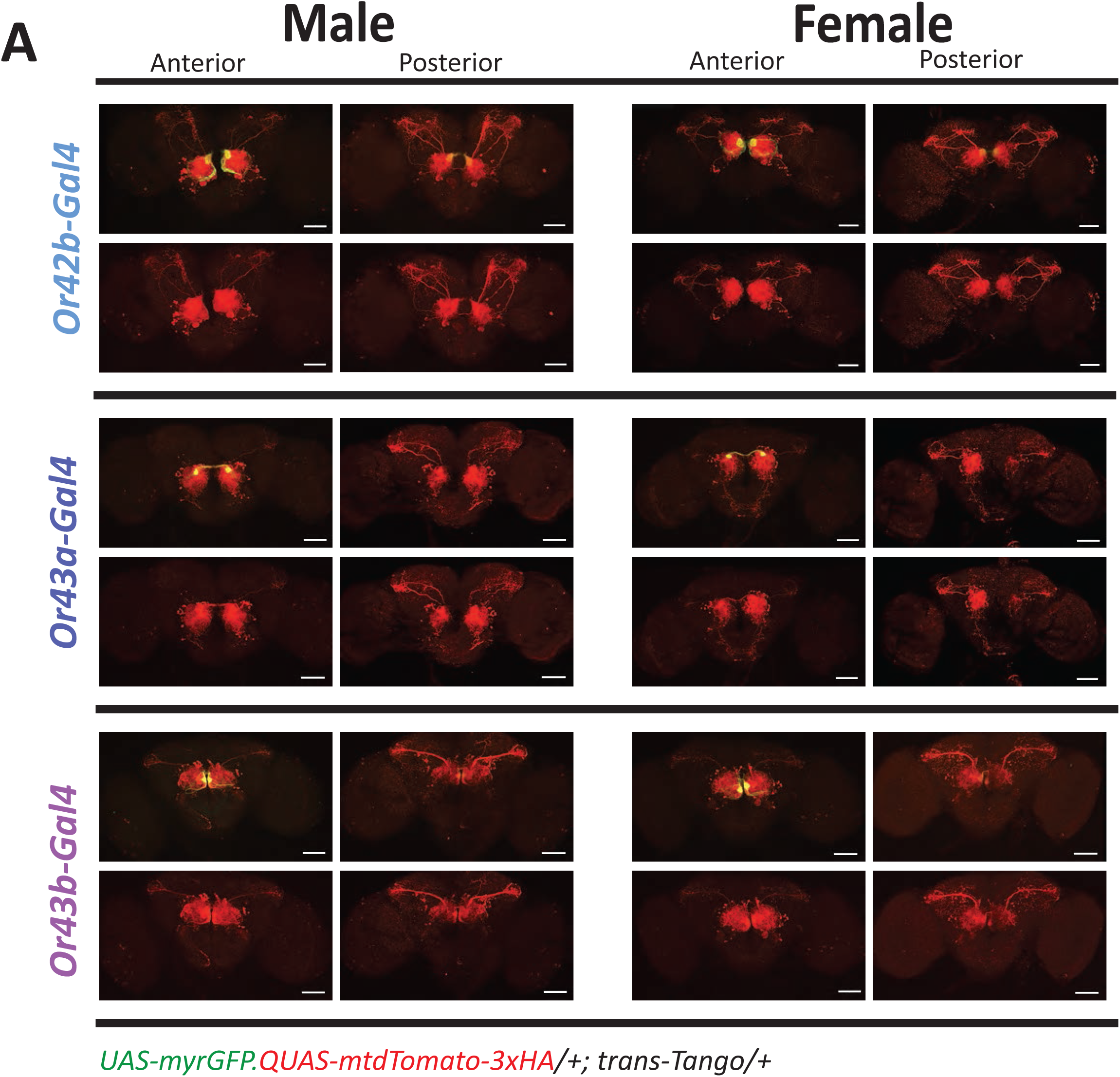
*trans*-TANGO identifies neuronal targets of *Or42b*, *Or43a*, and *Or43b*-expressing neurons and the brain regions they innervate. (A) Confocal maximum intensity projections of brains from 14–19-day-old male and female adult flies, aged at 18°C post-eclosion. The *trans*-TANGO reporter system enables visualization of odorant receptor-expressing neurons (green) and their postsynaptic targets (red). Flies carry the following transgenes: *UAS-myrGFP, QUAS-mtdTomato-3xHA; trans-TANGO; Or-Gal4*. The QF transcription factor is expressed in the postsynaptic target neurons of *Or-Gal4*-expressing cells. Scale bars: 50 μm.

The second-order projection neurons (PN) innervated by *Or42b* olfactory neurons project to the protocerebrum and terminate dorsally near the mushroom body calyx and lateral horn. In the EM micrograph they appear to be cell types: DM1_lPN and M_l2PNl20. We also observe additional projections in the top part of the suboesophageal ganglion (SEZ). The *Or42b* PN neuron cell bodies mainly reside lateral to the antennal lobe.

Similarly, *Or43a* PNs also project to protocerebrum and terminate dorsally near the mushroom body calyx and lateral horn (**Figure 10**). While both *Or42b* and *Or43a* PNs have two axon bundles projecting to the protocerebrum, *Or43a* both terminate more laterally. In the EM micrograph they appear to be cell type: DA4l_adPN. In addition, in females, the SEZ projections for *Or43a* PNs cover a broader area than those observed for *Or42b* PNs, meeting in the posterior SEZ region. The *Or43a* PN cell bodies are also located lateral to the antennal lobe.

In contrast, the *Or43b* PNs has one main axon bundle that terminates in one region of the lateral protocerebrum near the dorsoposterior edge of the lateral horn (**Figure 10**). Interestingly, *Or67d* PNs, which are involved in male pheromone detection, also terminate in this region of the lateral protocerebrum, near the lateral horn. In the EM micrograph the *Or43b* PNs appear to be cell type: VM2_adPN. The cell bodies for the *Or43b* PNs reside more anteriorly than observed for *Or42b* and *Or43a* PNs.

Previous studies have shown that odors are represented in different regions of the lateral horn and mushroom body, with fruit odors and pheromone odors occupying some overlapping domains (JEFFERIS *et al*. 2007; STRUTZ *et al*. 2014; BATES *et al*. 2020; CHOI *et al*. 2022). Furthermore, at least one of the receptors, *Or42b* has also been shown to be involved in food detection. The PNs for *Or42b*, *Or43a* and *Or43b* innervate several higher order processing regions, suggesting that detection of conspecific and heterospecific mates involves integration of sensory information across multiple brain regions, as well as integrating other odor information, including food odors.

## Discussion

This study provides an analysis of courtship behavior in *D. melanogaster*, *D. simulans*, and their hybrids, to gain insight into the molecular basis for species differences in behaviors, which will also reveal general principles regarding how behaviors are specified by the genome. The study integrates behavioral assays, neuroanatomical mapping, and molecular profiling to uncover mechanisms underlying species differences. Hybrid males exhibited a unique combination of courtship behaviors drawn from both parental species, some of which have been previously described (VON SCHILCHER AND MANNING 1975; WOOD AND RINGO 1980; WOOD *et al*. 1980; SHAHANDEH *et al*. 2020). Notably here, their behavioral output was modulated by the identity of the female target, with distinct patterns of wing extension, scissoring, and abdomen bobbing depending on whether the female was *D. melanogaster*, *D. simulans*, or hybrid. This target-dependent plasticity suggests that hybrid males possess a flexible behavioral repertoire, capable of tuning courtship strategies based on sensory input. In hybrids, courtship is not a rigid sequence; instead, it appears that behavioral modules can be selectively activated. This provides a framework for thinking about how behavioral elements can be added or subtracted within a species over evolutionary time. In addition, this suggests that additional insights into *D. melanogaster* courtship behavior may be gleaned from examining the impact of female phenotypes or “attractiveness” on male behavior, an area of research that is relatively underexplored.

Given that *D. melanogaster* and *D. simulans* are evolutionarily related, so is the neural circuitry underlying their behaviors. The genome provides the directions for circuit assembly by assigning the fate of neurons and directing which neurons will be connected. Mutations can affect these processes, through transcription factor expression modifications or expression changes in effector genes, making different aspects of circuit function a substrate for behavioral evolution (TOSCHES 2017). Variation in physiology (e.g. neurotransmitters, receptors, ion channels), connectivity (e.g. adhesion molecules), and developmental assignment (e.g. transcription factors) can result in circuits with different properties (Arendt et al. 2016, Hobert 2011), with an example of this already found with species-specific 7,11-HD pheromone preferences for *Drosophila* courtship behavior (SEEHOLZER *et al*. 2018). Despite behavioral differences of the hybrids compared to *D. melanogaster*, the neuroanatomical architecture of *fru P1*-expressing neurons was largely conserved. Using genetic intersectional tools and image registration, we observed similar spatial patterns and levels of individual variation across groups. This suggests that behavioral divergence may arise not from gross anatomical changes, but from differences in gene regulation or neuronal activity within a shared structural framework. This is consistent with previous results that demonstrated that the *fru* locus is not the evolutionary substrate responsible for behavioral differences across species (CANDE *et al*. 2014).

Our preference assays revealed that visual and chemical cues both contribute to species recognition, with eye color playing an important role. These results underscore the importance of multimodal cue integration in mate discrimination and suggest that hybrids may be particularly sensitive to cue conflict or ambiguity. We found that the genetic background of the *D. melanogaster* female parent significantly influenced hybrid male behavior. Hybrid_2 males, derived from a different maternal strain than Hybrid_1, showed reduced courtship toward *D. melanogaster* females and a stronger preference for *D. simulans* females. This highlights the role of genetic background in shaping behavioral phenotypes, particularly in hybrid contexts where regulatory interactions may be less canalized than in pure species.

The CUT&Tag analysis identified thousands of Fru^M^ target genes in *D. melanogaster* and *D. simulans*, with a substantial number being species-specific. While shared targets were enriched for known Fru DNA-binding motifs (DALTON *et al*. 2013), species-specific targets were depleted for these motifs, suggesting alternative regulatory mechanisms. These may include binding to novel motifs or indirect regulation via species-specific cofactors. The lack of overlap between species-specific targets and single-cell marker genes further suggests that these targets may act in broader sets of neurons, thus precluding their identification as marker genes (PALMATEER *et al*. 2023). We note that given that we mapped all the CUT&Tag sequence data to the *D. melanogaster* genome, to take advantage of the more complete annotation, we could be missing *D. simulans*-specific target genes that are not present in the *D. melanogaster* genome. Despite this limitation, more Fru^M^ target genes were identified in the *D. simulans* datasets, underscoring the robustness of the approach. Further examination of the data with mapping to the *D. simulans* genome may yield additional species-specific targets. Our previous work showed larger sex differences in gene expression in *fru P1*-expressing during pupal stages (PALMATEER *et al*. 2021), so identifying species-specific targets at earlier stages will also be important in the future, especially in determining differences in how the potential for behavior is specified developmentally. For this study we focused on the role *Or/Gr* genes that are bound by Fru^M^ specifically in *D. melanogaster*, using more labor-intensive behavioral screens. There are many additional candidates identified in this study that may have a role in reproductive isolation, including the set of genes that encode “Ion Channels”, given their known roles in sensory functions, especially detecting pheromones (LU *et al*. 2012; THISTLE *et al*. 2012a; TODA *et al*. 2012; LU *et al*. 2014). We provide the full list of Fru^M^ target genes, to facilitate further exploration.

While the *Or/Gr* genes are some of the best studied genes in *D. melanogaster*, this study reveals that additional screens will uncover new functional roles. Here, a genetic screen targeting *Or/Gr ∩ fru P1* neurons identified specific chemosensory circuits that influence mate discrimination. Silencing neurons expressing *Or42b*, *Or43a*, and *Or43b* disrupted copulation preferences, increasing copulation attempts toward *D. simulans* females. These findings implicate specific olfactory pathways in species recognition and provide candidate circuits for further functional dissection. Given the second order neurons downstream of neurons that express *Or42b*, *Or43a*, and *Or43b* project to different regions of the lateral horn and mushroom body, reveals that the information is integrated by distinct higher order processing centers.

Furthermore, *Or43a* is the only one of the three to have projections in females to the suboesophageal ganglion, which could underlie sex differences in olfaction important for reproductive behaviors. *Or42b*-expressing neurons have previously been implicated in food detection (SEMMELHACK AND WANG 2009; ROOT *et al*. 2011), consistent with the idea that mating behaviors are linked to food substrates (ANHOLT *et al*. 2020). These neuronal subsets provide an ideal system to understand how distinct types of information are integrated across higher order processing centers to bring about a behavior.

Understanding how rapidly evolving species-specific courtship behaviors are genetically specified offers critical insight into the broader mechanisms that govern behavioral specification. While this study focused on *fru,* the sex hierarchy gene *doublesex* is also a key regulator of species-specific courtship behavior (YE *et al*. 2024). Elucidating how *fru* and *dsx* coordinate to direct evolutionary changes is an important future goal. We expect that the further examination of Fru^M^ and Dsx targets in other species will be highly informative in understanding how behavioral changes arise over evolutionary time, especially identifying species-specific Fru^M^ and Dsx targets during development. Integrating genomic approaches coupled with functional studies is a powerful strategy for uncovering the mechanisms by which behavioral elements are modified across evolutionary timescales.

## Data availability statement

All data necessary for confirming the conclusions are presented in the manuscript. The CUT&Tag data will be available at the GEO repository.

## Acknowledgements

We thank the Bloomington Drosophila Stock Center (NIH P40OD018537) and Drosophila colleagues for generously providing stocks and antibodies. Several antibodies used in this study were obtained from the Developmental Studies Hybridoma Bank, created by the NICHD of the NIH, and maintained at The University of Iowa, Department of Biology, Iowa City, IA 52242. We used FlyBase to find information about genes, stocks, phenotypes, and function (OZTURK-COLAK *et al*. 2024). We thank Adela Peña for contributions to some of the experiments presented. Microsoft Co-Pilot was used to copy edit the full document. For some of our figure schematics, we used Biorender (citation: *Created in BioRender. Arbeitman, M. (2025)* https://BioRender.com/v3me7zc).

---

**Video 1. Hybrid_1 males courting *D. melanogaster* and *D. simulans* females.**

This video shows Hybrid_1 males courting w; Berlin (*D. melanogaster*) and Dsim167.4 (*D. simulans*) females. The male behavior changes depending on the species he is courting.

## References

Ahmad, K., 2020 CUT&Tag with Drosophila tissues. protocols.io.

Aleksander, S. A., J. Balhoff, S. Carbon, J. M. Cherry, H. J. Drabkin et al., 2023 The Gene Ontology knowledgebase in 2023. Genetics 224.

Anholt, R. R. H., P. O’Grady, M. F. Wolfner and S. T. Harbison, 2020 Evolution of Reproductive Behavior. Genetics 214: 49–73.

Ashburner, M., C. A. Ball, J. A. Blake, D. Botstein, H. Butler et al., 2000 Gene Ontology: tool for the unification of biology. Nature Genetics 25: 25–29.

Auer, T., and R. Benton, 2016 Sexual circuitry in Drosophila. Current Opinion in Neurobiology 38: 18–26.

Averhoff, W. W., and R. H. Richardson, 1974 Pheromonal Control of Mating Patterns in Drosophila-Melanogaster. Behavior Genetics 4: 207–225.

Baker, C. A., X. J. Guan, M. Choi and M. Murthy, 2024 The role of fruitless in specifying courtship behaviors across divergent Drosophila species. Sci Adv 10: eadk1273.

Barbash, D. A., 2010 Ninety years of Drosophila melanogaster hybrids. Genetics 186: 1–8.

Barbash, D. A., D. F. Siino, A. M. Tarone and J. Roote, 2003 A rapidly evolving MYB-related protein causes species isolation in Drosophila. Proceedings of the National Academy of Sciences of the United States of America 100: 5302–5307.

Bates, A. S., P. Schlegel, R. J. V. Roberts, N. Drummond, I. F. M. Tamimi et al., 2020 Complete Connectomic Reconstruction of Olfactory Projection Neurons in the Fly Brain. Curr Biol 30: 3183–3199 e3186.

Billeter, J. C., J. Atallah, J. J. Krupp, J. G. Millar and J. D. Levine, 2009 Specialized cells tag sexual and species identity in Drosophila melanogaster. Nature 461: 987–U250.

Bontonou, G., and C. Wicker-Thomas, 2014 Sexual Communication in the Drosophila Genus. Insects 5: 439–458.

Brideau, N. J., H. A. Flores, J. Wang, S. Maheshwari, X. Wang et al., 2006 Two Dobzhansky-Muller genes interact to cause hybrid lethality in Drosophila. Science 314: 1292–1295.

Brovero, S. G., J. C. Fortier, H. Hu, P. C. Lovejoy, N. R. Newell et al., 2021 Investigation of Drosophilafruitless neurons that express Dpr/DIP cell adhesion molecules. eLife 10.

Brovkina, M. V., R. Duffie, A. E. C. Burtis and E. J. Clowney, 2021 Fruitless decommissions regulatory elements to implement cell-type-specific neuronal masculinization. Plos Genetics 17.

Bushnell, B., J. Rood and E. Singer, 2017 BBMerge - Accurate paired shotgun read merging via overlap. Plos One 12.

Cachero, S., A. D. Ostrovsky, J. Y. Yu, B. J. Dickson and G. Jefferis, 2010 Sexual Dimorphism in the Fly Brain. Current Biology 20: 1589–1601.

Cande, J., P. Andolfatto, B. Prud’homme, D. L. Stern and N. Gompel, 2012 Evolution of Multiple Additive Loci Caused Divergence between Drosophila yakuba and D-santomea in Wing Rowing during Male Courtship. Plos One 7.

Cande, J., D. L. Stern, T. Morita, B. Prud’homme and N. Gompel, 2014 Looking under the lamp post: neither fruitless nor doublesex has evolved to generate divergent male courtship in Drosophila. Cell Rep 8: 363–370.

Cattani, M. V., and D. C. Presgraves, 2009 Genetics and Lineage-Specific Evolution of a Lethal Hybrid Incompatibility Between Drosophila mauritiana and Its Sibling Species. Genetics 181: 1545–1555.

Cattani, M. V., and D. C. Presgraves, 2012 Incompatibility between X chromosome factor and pericentric heterochromatic region causes lethality in hybrids between Drosophila melanogaster and its sibling species. Genetics 191: 549–559.

Chatterjee, R. N., P. Chatterjee, A. Pal and M. Pal-Bhadra, 2007 Drosophila simulans Lethal hybrid rescue mutation (Lhr) rescues inviable hybrids by restoring X chromosomal dosage compensation and causes fluctuating asymmetry of development. Journal of Genetics 86: 203–215.

Choi, K., W. K. Kim and C. Hyeon, 2022 Olfactory responses of Drosophila are encoded in the organization of projection neurons. Elife 11.

Clark, A. G., M. B. Eisen, D. R. Smith, C. M. Bergman, B. Oliver et al., 2007 Evolution of genes and genomes on the phylogeny. Nature 450: 203–218.

Clowney, E. J., S. Iguchi, J. J. Bussell, E. Scheer and V. Ruta, 2015 Multimodal Chemosensory Circuits Controlling Male Courtship in Drosophila. Neuron 87: 1036–1049.

Clynen, E., L. Ciudad, X. Bellés and M. D. Piulachs, 2011 Conservation of fruitless’ role as master regulator of male courtship behaviour from cockroaches to flies. Development Genes and Evolution 221: 43–48.

Cobb, M., B. Burnet and K. Connolly, 1986 THE STRUCTURE OF COURTSHIP IN THE DROSOPHILA-MELANOGASTER SPECIES SUBGROUP. Behaviour 97: 182–212.

Cobb, M., K. Connolly and B. Burnet, 1985 COURTSHIP BEHAVIOR IN THE MELANOGASTER SPECIES SUBGROUP OF DROSOPHILA, pp. 203–231.

Coleman, R. T., I. Morantte, G. T. Koreman, M. L. Cheng, Y. Ding et al., 2024 A modular circuit coordinates the diversification of courtship strategies. Nature 635: 142–150.

Combs, P. A., J. J. Krupp, N. M. Khosla, D. Bua, D. A. Petrov et al., 2018 Tissue-Specific -Regulatory Divergence Implicates in Inhibiting Interspecies Mating in. Current Biology 28: 3969-+.

Cooper, J. C., A. Lukacs, S. Reich, T. Schauer, A. Imhof et al., 2019 Altered Localization of Hybrid Incompatibility Proteins in. Molecular Biology and Evolution 36: 1783–1792.

Couto, A., M. Alenius and B. J. Dickson, 2005 Molecular, anatomical, and functional organization of the Drosophila olfactory system. Curr Biol 15: 1535–1547.

Coyne, J. A., and H. A. Orr, 2004 Speciation. Speciation.: i-xiii, 1-545.

Cuykendall, T. N., P. Satyaki, S. Ji, D. M. Clay, N. B. Edelman et al., 2014 A screen for F1 hybrid male rescue reveals no major-effect hybrid lethality loci in the Drosophila melanogaster autosomal genome. G3 (Bethesda) 4: 2451-2460.

Dalton, J. E., J. M. Fear, S. Knott, B. S. Baker, L. M. McIntyre et al., 2013 Male-specific Fruitless isoforms have different regulatory roles conferred by distinct zinc finger DNA binding domains. Bmc Genomics 14.

Dalton, J. E., M. S. Lebo, L. E. Sanders, F. Sun and M. N. Arbeitman, 2009 Ecdysone receptor acts in fruitless-expressing neurons to mediate drosophila courtship behaviors. Curr Biol 19: 1447–1452.

Danecek, P., J. K. Bonfield, J. Liddle, J. Marshall, V. Ohan et al., 2021 Twelve years of SAMtools and BCFtools. Gigascience 10.

David, J. R., F. Lemeunier, L. Tsacas and A. Yassin, 2007 The historical discovery of the nine species in the Drosophila melanogaster species subgroup. Genetics 177: 1969–1973.

Dunipace, L., S. Meister, C. McNealy and H. Amrein, 2001 Spatially restricted expression of candidate taste receptors in the Drosophila gustatory system. Curr Biol 11: 822–835.

Dweck, H. K., S. A. Ebrahim, M. Thoma, A. A. Mohamed, I. W. Keesey et al., 2015 Pheromones mediating copulation and attraction in Drosophila. Proc Natl Acad Sci U S A 112: E2829–2835.

Fabre, C. C., B. Hedwig, G. Conduit, P. A. Lawrence, S. F. Goodwin et al., 2012 Substrate-borne vibratory communication during courtship in Drosophila melanogaster. Curr Biol 22: 2180–2185.

Fan, P., D. S. Manoli, O. M. Ahmed, Y. Chen, N. Agarwal et al., 2013 Genetic and Neural Mechanisms that Inhibit Drosophila from Mating with Other Species. Cell 154: 89–102.

Ferveur, J. F., 1997 The pheromonal role of cuticular hydrocarbons in Drosophila melanogaster. Bioessays 19: 353–358.

Ferveur, J. F., and G. Sureau, 1996 Simultaneous influence on male courtship of stimulatory and inhibitory pheromones produced by live sex-mosaic Drosophila melanogaster. Proceedings of the Royal Society B-Biological Sciences 263: 967–973.

Fishilevich, E., A. I. Domingos, K. Asahina, F. Naef, L. B. Vosshall et al., 2005 Chemotaxis behavior mediated by single larval olfactory neurons in Drosophila. Curr Biol 15: 2086–2096.

Fishilevich, E., and L. B. Vosshall, 2005 Genetic and functional subdivision of the Drosophila antennal lobe. Curr Biol 15: 1548–1553.

Goldman, T. D., and M. N. Arbeitman, 2007 Genomic and functional studies of Drosophila sex hierarchy regulated gene expression in adult head and nervous system tissues. PLoS Genet 3: e216.

Greenspan, R. J., and J. F. Ferveur, 2000 Courtship in Drosophila. Annual Review of Genetics 34: 205–232.

Henikoff, S., J. G. Henikoff, H. S. Kaya-Okur and K. Ahmad, 2020 Efficient chromatin accessibility mapping in situ by nucleosome-tethered tagmentation. Elife 9.

Hodge, J. J., 2009 Ion channels to inactivate neurons in Drosophila. Front Mol Neurosci 2: 13.

Hollocher, H., and C. I. Wu, 1996 The genetics of reproductive isolation in the Drosophila simulans clade: X vs autosomal effects and male vs female effects. Genetics 143: 1243–1255.

Hu, Y. H., A. Comjean, H. Attrill, G. Antonazzo, J. Thurmond et al., 2023 PANGEA: a new gene set enrichment tool for and common research organisms. Nucleic Acids Research 51: W419–W426.

Huang, W., R. Loganantharaj, B. Schroeder, D. Fargo and L. Li, 2013 PAVIS: a tool for Peak Annotation and Visualization. Bioinformatics 29: 3097–3099.

Inagaki, H. K., Y. Jung, E. D. Hoopfer, A. M. Wong, N. Mishra et al., 2014 Optogenetic control of Drosophila using a red-shifted channelrhodopsin reveals experience-dependent influences on courtship. Nature Methods 11: 325–U311.

Ishii, K., M. Wohl, A. DeSouza and K. Asahina, 2020 Sex-determining genes distinctly regulate courtship capability and target preference via sexually dimorphic neurons. Elife 9.

Ito, H., K. Sato, M. Koganezawa, M. Ote, K. Matsumoto et al., 2012 Fruitless Recruits Two Antagonistic Chromatin Factors to Establish Single-Neuron Sexual Dimorphism. Cell 149: 1327–1338.

Jallon, J. M., 1984 A FEW CHEMICAL WORDS EXCHANGED BY DROSOPHILA DURING COURTSHIP AND MATING. Behavior Genetics 14: 441–478.

Jallon, J. M., and J. R. David, 1987 VARIATIONS IN CUTICULAR HYDROCARBONS AMONG THE 8 SPECIES OF THE DROSOPHILA-MELANOGASTER SUBGROUP. Evolution 41: 294–302.

Jefferis, G. S. X. E., C. J. Potter, A. I. Chan, E. C. Marin, T. Rohlfing et al., 2007 Comprehensive maps of higher offactory centers:: Spatially segregated fruit and pheromone representation. Cell 128: 1187–1203.

Jezovit, J. A., J. D. Levine and J. Schneider, 2017 Phylogeny, environment and sexual communication across the Drosophila genus. Journal of Experimental Biology 220: 42–52.

Kallman, B. R., H. Kim and K. Scott, 2015 Excitation and inhibition onto central courtship neurons biases Drosophila mate choice. Elife 4.

Kaya-Okur, H. S., S. J. Wu, C. A. Codomo, E. S. Pledger, T. D. Bryson et al., 2019 CUT&Tag for efficient epigenomic profiling of small samples and single cells. Nat Commun 10: 1930.

Kimura, K. I., M. Ote, T. Tazawa and D. Yamamoto, 2005 Fruitless specifies sexually dimorphic neural circuitry in the Drosophila brain. Nature 438: 229–233.

Koganezawa, M., D. Haba, T. Matsuo and D. Yamamoto, 2010 The Shaping of Male Courtship Posture by Lateralized Gustatory Inputs to Male-Specific Interneurons. Current Biology 20: 1–8.

Kohatsu, S., M. Koganezawa and D. Yamamoto, 2011 Female Contact Activates Male-Specific Interneurons that Trigger Stereotypic Courtship Behavior in Drosophila. Neuron 69: 498–508.

Kurtovic, A., A. Widmer and B. J. Dickson, 2007 A single class of olfactory neurons mediates behavioural responses to a Drosophila sex pheromone. Nature 446: 542–546.

Kwon, J. Y., A. Dahanukar, L. A. Weiss and J. R. Carlson, 2011 Molecular and cellular organization of the taste system in the Drosophila larva. J Neurosci 31: 15300–15309.

Langmead, B., and S. L. Salzberg, 2012 Fast gapped-read alignment with Bowtie 2. Nature Methods 9: 357–U354.

Lebo, M. S., L. E. Sanders, F. Sun and M. N. Arbeitman, 2009 Somatic, germline and sex hierarchy regulated gene expression during Drosophila metamorphosis. BMC Genomics 10: 80.

Lebreton, S., F. Borrero-Echeverry, F. Gonzalez, M. Solum, E. A. Wallin et al., 2017 A Drosophila female pheromone elicits species-specific long-range attraction via an olfactory channel with dual specificity for sex and food. BMC Biol 15: 88.

Lee, W. H., and T. K. Watanabe, 1987 EVOLUTIONARY GENETICS OF THE DROSOPHILA-MELANOGASTER SUBGROUP. 1. PHYLOGENETIC-RELATIONSHIPS BASED ON MATINGS, HYBRIDS AND PROTEINS. Japanese Journal of Genetics 62: 225-239.

Lu, B., A. LaMora, Y. Sun, M. J. Welsh and Y. Ben-Shahar, 2012 ppk23-Dependent Chemosensory Functions Contribute to Courtship Behavior in Drosophila melanogaster. Plos Genetics 8.

Lu, B., K. M. Zelle, R. Seltzer, A. Hefetz and Y. Ben-Shahar, 2014 Feminization of pheromone-sensing neurons affects mating decisions in Drosophila males. Biology Open 3: 152–160.

Luo, Y., G. J. S. Talross and J. R. Carlson, 2024 Function and evolution of Ir52 receptors in mate detection in Drosophila. Curr Biol 34: 5395–5408 e5396.

Lyne, R., R. Smith, K. Rutherford, M. Wakeling, A. Varley et al., 2007 FlyMine: an integrated database for Drosophila and Anopheles genomics. Genome Biol 8: R129.

Manning, A., 1959 The sexual isolation between D*rosophila melanogaster* and D*rosophila simulans*. Volume 7: 60–65.

Manoli, D. S., M. Foss, A. Villella, B. J. Taylor, J. C. Hall et al., 2005 Male-specific fruitless specifies the neural substrates of Drosophila courtship behaviour. Nature 436: 395–400.

Masly, J. P., C. D. Jones, M. A. F. Noor, J. Locke and H. A. Orr, 2006 Gene transposition as a cause of hybrid sterility in Drosophila. Science 313: 1448–1450. Matsliah, Codex: Connectome Data Explorer.

McKinney, R. M., and Y. Ben-Shahar, 2019 Visual recognition of the female body axis drives spatial elements of male courtship in *Drosophila melanogaster*. bioRxiv: 576322.

McLaughlin, C. N., M. Brbic, Q. Xie, T. Li, F. Horns et al., 2021 Single-cell transcriptomes of developing and adult olfactory receptor neurons in Drosophila. Elife 10.

Meers, M. P., D. Tenenbaum and S. Henikoff, 2019 Peak calling by Sparse Enrichment Analysis for CUT&RUN chromatin profiling. Epigenetics & Chromatin 12.

Neville, M. C., T. Nojima, E. Ashley, D. J. Parker, J. Walker et al., 2014 Male-Specific Fruitless Isoforms Target Neurodevelopmental Genes to Specify a Sexually Dimorphic Nervous System. Current Biology 24: 229–241.

Ostrovsky, A., S. Cachero and G. Jefferis, 2013 Clonal analysis of olfaction in Drosophila: immunochemistry and imaging of fly brains. Cold Spring Harbor protocols 2013: 342–346.

Ozturk-Colak, A., S. J. Marygold, G. Antonazzo, H. Attrill, D. Goutte-Gattat et al., 2024 FlyBase: updates to the Drosophila genes and genomes database. Genetics 227.

Palmateer, C. M., C. Artikis, S. G. Brovero, B. Friedman, A. Gresham et al., 2023 Single-cell transcriptome profiles of Drosophila fruitless-expressing neurons from both sexes. Elife 12.

Palmateer, C. M., S. C. Moseley, S. Ray, S. G. Brovero and M. N. Arbeitman, 2021 Analysis of cell-type-specific chromatin modifications and gene expression in Drosophila neurons that direct reproductive behavior. PLoS Genet 17: e1009240.

Pan, Y., G. W. Meissner and B. S. Baker, 2012 Joint control of Drosophila male courtship behavior by motion cues and activation of male-specific P1 neurons. Proceedings of the National Academy of Sciences of the United States of America 109: 10065–10070.

Pardy, J. A., H. D. Rundle, M. A. Bernards and A. J. Moehring, 2019 The genetic basis of female pheromone differences between Drosophila melanogaster and D-simulans. Heredity 122: 93–109.

Parker, D. J., A. Gardiner, M. C. Neville, M. G. Ritchie and S. F. Goodwin, 2014 The evolution of novelty in conserved genes; evidence of positive selection in the gene is localised to alternatively spliced exons. Heredity 112: 300–306.

Peng, Q., J. Chen and Y. Pan, 2021 From fruitless to sex: On the generation and diversification of an innate behavior. Genes Brain Behav 20: e12772.

Phadnis, N., E. P. Baker, J. C. Cooper, K. A. Frizzell, E. Hsieh et al., 2015 An essential cell cycle regulation gene causes hybrid inviability in Drosophila. Science 350: 1552–1555.

Presgraves, D. C., L. Balagopalan, S. M. Abmayr and H. A. Orr, 2003 Adaptive evolution drives divergence of a hybrid inviability gene between two species of Drosophila. Nature 423: 715–719.

Presgraves, D. C., and J. P. Masly, 2007 High-Resolution Genome-Wide Dissection of the Two Rules of Speciation in Drosophila. Figshare.

Rohlfing, T., 2012 Image Similarity and Tissue Overlaps as Surrogates for Image Registration Accuracy: Widely Used but Unreliable. Ieee Transactions on Medical Imaging 31: 153–163.

Rohlfing, T., and C. R. Maurer, 2003 Nonrigid image registration in shared-memory multiprocessor environments with application to brains, breasts, and bees. Ieee Transactions on Information Technology in Biomedicine 7: 16–25.

Root, C. M., K. I. Ko, A. Jafari and J. W. Wang, 2011 Presynaptic facilitation by neuropeptide signaling mediates odor-driven food search. Cell 145: 133–144.

Sanders, L. E., and M. N. Arbeitman, 2008 Doublesex establishes sexual dimorphism in the Drosophila central nervous system in an isoform-dependent manner by directing cell number. Developmental Biology 320: 378–390.

Sato, K., R. Tanaka, Y. Ishikawa and D. Yamamoto, 2020 Behavioral Evolution of Drosophila: Unraveling the Circuit Basis. Genes (Basel) 11.

Sato, K., and D. Yamamoto, 2020 Contact-Chemosensory Evolution Underlying Reproductive Isolation in Drosophila Species. Frontiers in Behavioral Neuroscience 14.

Sato, K., and D. Yamamoto, 2023 Molecular and cellular origins of behavioral sex differences: a tiny little fly tells a lot. Front Mol Neurosci 16: 1284367.

Sawamura, K., and M. T. Yamamoto, 1997 Characterization of a reproductive isolation gene, zygotic hybrid rescue, of Drosophila melanogaster by using minichromosomes. Heredity 79: 97–103.

Seeholzer, L. F., M. Seppo, D. L. Stern and V. Ruta, 2018 Evolution of a central neural circuit underlies Drosophila mate preferences. Nature 559: 564-+.

Semmelhack, J. L., and J. W. Wang, 2009 Select Drosophila glomeruli mediate innate olfactory attraction and aversion. Nature 459: 218–223.

Shahandeh, M. P., A. Pischedda, J. M. Rodriguez and T. L. Turner, 2020 The Genetics of Male Pheromone Preference Difference Between Drosophila melanogaster and Drosophila simulans. G3-Genes Genomes Genetics 10: 401-415.

Shahandeh, M. P., A. Pischedda and T. L. Turner, 2018 Male mate choice via cuticular hydrocarbon pheromones drives reproductive isolation between Drosophila species. Evolution 72: 123–135.

Shirangi, T. R., H. D. Dufour, T. M. Williams and S. B. Carroll, 2009 Rapid Evolution of Sex Pheromone-Producing Enzyme Expression in Drosophila. Plos Biology 7.

Spieth, H. T., 1952 Mating behavior within the genus Drosophila (Diptera). American Museum of Natural History.

Spieth, H. T., 1974 Courtship behavior in Drosophila. Annu Rev Entomol 19: 385–405.

Stockinger, P., D. Kvitsiani, S. Rotkopf, L. Tirian and B. J. Dickson, 2005 Neural circuitry that governs Drosophila male courtship behavior. Cell 121: 795–807.

Strutz, A., J. Soelter, A. Baschwitz, A. Farhan, V. Grabe et al., 2014 Decoding odor quality and intensity in the Drosophila brain. Elife 3: e04147.

Sturtevant, A. H., 1915 Experiments in sexual recognition and the problems of sexual selection in Drosophila.

Sturtevant, A. H., 1920 Genetic studies on drosophila simulans. I. Introduction. Hybrids with Drosophila melanogaster. Genetics 5: 488–500.

Sturtevant, A. H., 1921 Genetic studies on Drosophila simulans. III. Autosomal genes. General discussion. Genetics 6: 179–207.

Talay, M., E. B. Richman, N. J. Snell, G. G. Hartmann, J. D. Fisher et al., 2017 Transsynaptic Mapping of Second-Order Taste Neurons in Flies by trans-Tango. Neuron 96: 783–795 e784.

Tamura, K., S. Subramanian and S. Kumar, 2004 Temporal patterns of fruit fly (Drosophila) evolution revealed by mutation clocks. Molecular Biology and Evolution 21: 36–44.

Tanaka, R., T. Higuchi, S. Kohatsu, K. Sato and D. Yamamoto, 2017 Optogenetic Activation of the fruitless-Labeled Circuitry in Drosophila subobscura Males Induces Mating Motor Acts. J Neurosci 37: 11662–11674.

Tang, S., and D. C. Presgraves, 2015 Lineage-Specific Evolution of the Complex Nup160 Hybrid Incompatibility Between Drosophila melanogaster and Its Sister Species. Genetics 200: 1245–1254.

Tao, Y., Z. B. Zeng, J. Li, D. L. Hartl and C. C. Laurie, 2003 Genetic dissection of hybrid Incompatibilities between Drosophila simulansand D. mauritiana. II. Mapping hybrid male sterility loci on the third chromosome. Genetics 164: 1399–1418.

Thistle, R., P. Cameron, A. Ghorayshi, L. Dennison and K. Scott, 2012a Contact Chemoreceptors Mediate Male-Male Repulsion and Male-Female Attraction during Drosophila Courtship. Cell 149: 1140–1151.

Thistle, R., P. Cameron, A. Ghorayshi, L. Dennison and K. Scott, 2012b Contact chemoreceptors mediate male-male repulsion and male-female attraction during Drosophila courtship. Cell 149: 1140–1151.

Ting, C. T., S. C. Tsaur, M. L. Wu and C. I. Wu, 1998 A rapidly evolving homeobox at the site of a hybrid sterility gene. Science 282: 1501–1504.

Toda, H., X. Zhao and B. J. Dickson, 2012 The Drosophila Female Aphrodisiac Pheromone Activates ppk23(+) Sensory Neurons to Elicit Male Courtship Behavior. Cell Reports 1: 599–607.

Tootoonian, S., P. Coen, R. Kawai and M. Murthy, 2012 Neural representations of courtship song in the Drosophila brain. J Neurosci 32: 787–798.

Tosches, M. A., 2017 Developmental and genetic mechanisms of neural circuit evolution. Dev Biol 431: 16–25.

True, J. R., J. M. Mercer and C. C. Laurie, 1996 Differences in crossover frequency and distribution among three sibling species of Drosophila. Genetics 142: 507–523.

Vernes, S. C., 2014 Genome wide identification of Fruitless targets suggests a role in upregulating genes important for neural circuit formation. Scientific Reports 4.

Vernier, C. L., N. Leitner, K. M. Zelle, M. Foltz, S. Dutton et al., 2023 A pleiotropic chemoreceptor facilitates the production and perception of mating pheromones. iScience 26: 105882.

von Philipsborn, A. C., T. X. Liu, J. Y. Yu, C. Masser, S. S. Bidaye et al., 2011 Neuronal Control of Drosophila Courtship Song. Neuron 69: 509–522.

Von Schilcher, F., and A. Manning, 1975 COURTSHIP SONG AND MATING SPEED IN HYBRIDS BETWEEN DROSOPHILA-MELANOGASTER AND DROSOPHILA-SIMULANS. Behavior Genetics 5: 395–404.

Watanabe, T. K., 1979 GENE THAT RESCUES THE LETHAL HYBRIDS BETWEEN DROSOPHILA-MELANOGASTER AND D-SIMULANS. Japanese Journal of Genetics 54: 325–331.

Wei, K. H. C., A. G. Clark and D. A. Barbash, 2014 Limited Gene Misregulation Is Exacerbated by Allele-Specific Upregulation in Lethal Hybrids between and. Molecular Biology and Evolution 31: 1767–1778.

Wood, D., and J. M. Ringo, 1980 MALE MATING DISCRIMINATION IN DROSOPHILA-MELANOGASTER, DROSOPHILA-SIMULANS AND THEIR HYBRIDS. Evolution 34: 320–329.

Wood, D., J. M. Ringo and L. L. Johnson, 1980 ANALYSIS OF COURTSHIP SEQUENCES OF THE HYBRIDS BETWEEN DROSOPHILA-MELANOGASTER AND DROSOPHILA-SIMULANS. Behavior Genetics 10: 459–466.

Ye, D., J. T. Walsh, I. P. Junker and Y. Ding, 2024 Changes in the cellular makeup of motor patterning circuits drive courtship song evolution in Drosophila. Curr Biol 34: 2319–2329 e2316.

Yu, J. Y., M. I. Kanai, E. Demir, G. Jefferis and B. J. Dickson, 2010 Cellular Organization of the Neural Circuit that Drives Drosophila Courtship Behavior. Current Biology 20: 1602–1614.

Zheng Y, A. K., Henikoff S, 2020 CUT&Tag Data Processing and Analysis Tutorial. protocols.io.

